# Switching an active site helix in dihydrofolate reductase reveals limits to sub-domain modularity

**DOI:** 10.1101/2021.06.18.448971

**Authors:** Victor Y. Zhao, Joao V. Rodrigues, Elena R. Lozovsky, Daniel L. Hartl, Eugene I. Shakhnovich

## Abstract

To what degree are individual structural elements within proteins modular such that similar structures from unrelated proteins can be interchanged? We study sub-domain modularity by creating 20 chimeras of an enzyme, *E. coli* dihydrofolate reductase (DHFR), in which a catalytically important, 10-residue α-helical sequence is replaced by α-helical sequences from a diverse set of proteins. The chimeras stably fold but have a range of diminished thermal stabilities and catalytic activities. Evolutionary coupling analysis indicates that the residues of this α-helix are under selection pressure to maintain catalytic activity in DHFR. We performed molecular dynamics simulations using replica exchange with solute-tempering. Chimeras with low catalytic activity exhibit non-helical conformations that block the binding site and disrupt the positioning of the catalytically essential residue D27. Simulation observables and *in vitro* measurements of thermal stability and substrate binding affinity are strongly correlated. Several *E. coli* strains with chromosomally integrated chimeric DHFRs can grow, with growth rates that follow predictions from a kinetic flux model that depends on the intracellular abundance and catalytic activity of DHFR. Our findings show that although α-helices are not universally substitutable, the molecular and fitness effects of modular segments can be predicted by the biophysical compatibility of the replacement segment.

**Statement of Significance:** α-helices are ubiquitous components of protein structure that exhibit a degree of independent folding behavior, making them plausible structural modules within proteins. Here, we assess the effects of switching the sequence of an α-helix in an essential enzyme for α-helical sequences from evolutionarily unrelated proteins. The resultant chimeric proteins can still fold but enzymatic activity, stability, and cellular growth rates are negatively affected. Computational investigations reveal how residues in an α-helix have been shaped by selection pressure to maintain catalytic activity and a specific, helical conformation of the protein. More broadly, we illustrate how molecular and fitness effects of switching protein segments depend on the protein and cellular context.

## Introduction

Modularity in biology is characterized by a portion of a larger biological unit evolving and functioning as a distinct unit (1). By their ability to fold independently and carry out specific functions either alone or as part of a larger polypeptide, protein domains demonstrate properties of modularity. The modularity of protein domains facilitates evolution by enabling their combination and shuffling to create multi-domain proteins with novel functions (2–4).

While the evidence that protein domains can behave as independent units is clear, less is known about modularity within domains (5). Although short peptide sequences are unlikely to have defined structures in isolation, insertions, deletions, and shuffling of short segments commonly occur in protein evolution (6–10). One suggestion of sub-domain modularity is the occurrence of similar substructures in proteins from across the protein universe, from contiguous segments to recurring tertiary interactions (11–17). This reuse of structure reflects both natural thermodynamic basins of stability as well as inheritance of ancestral, peptide building blocks in evolution (15, 16, 18–20).

At the level of secondary structure, there is evidence that α-helices within proteins can act as distinct units with localized folding and unfolding transitions (21–24), and some α-helices exist as standalone structures (25–27). Compared to other structures in proteins, α-helices are also more tolerant of mutations (28, 29). A few past studies examined the replacement of whole α-helices in enzymes (30–32). In *E. coli* alkaline phosphatase, α-helices were replaced with non-homologous helical sequences while maintaining enzyme activity in most cases (30). More recently, we replaced the catalytically important α_B_-helix in *E. coli* dihydrofolate reductase (DHFR) with orthologous sequences from DHFRs of six other species with little effect on catalytic activity and organism growth rate but some changes to thermal stability and antibiotic resistance (32). For these reasons, we hypothesize that α-helices represent modular segments within protein domains.

In this work, we explore the modularity of α-helices using *E. coli* DHFR as a model protein. We performed non-homologous replacement by switching the α_B_-helix sequence in *E. coli* DHFR with sequences taken from helices in unrelated, non-DHFR proteins while leaving the key catalytic residue, D27, unchanged. The chimeras vary in catalytic activity, although all of them have reduced activity compared with wildtype DHFR. The chimeras with weaker catalytic activity have residues that appear infrequently among homologous DHFR sequences and exhibit non-helical conformations. The ability of replacement helix sequences to maintain a wildtype-like helical conformation determines cellular fitness through the dual effect of helix conformation on catalytic activity and cellular protein abundance, which is determined by the balance between rates of protein production and degradation (33). These findings support qualified modularity in this specific α-helix in which tertiary interactions govern the functional compatibility of replacement sequences. More generally, our work shows, in mechanistic detail, how structural variation in the vicinity of the active site broadly mediates the activity of an enzyme. On a cellular level, we show how large-scale sequence variation pleiotropically affects the fitness landscape through its dual effect on enzymatic activity and in-cell turnover, which cause pronounced variation in total DHFR activity in the cytoplasm.

## Materials and Methods

### Chimera generation process

DHFR is a globular protein consisting of a β-sheet surrounded by loops and helices (Fig. 1A). The protein structure can be divided into two subdomains: the adenosine-binding subdomain (residues 38-88, colored gray) and the major subdomain (residues 1-37 and 89-159, various colors). The major subdomain consists of a central β-sheet along with two of the three α-helices in DHFR as well as three loops: the Met 20 loop (residues 9-24), the FG loop (residues 116-132), and the GH loop (residues 142-150) (36). The 10 residues of the α_B_-helix targeted for replacement are highlighted in yellow. We targeted the α_B_-helix because of its position at the enzyme active site, where it positions catalytically essential D27 (37), shown in stick representation.

**Fig. 1:**
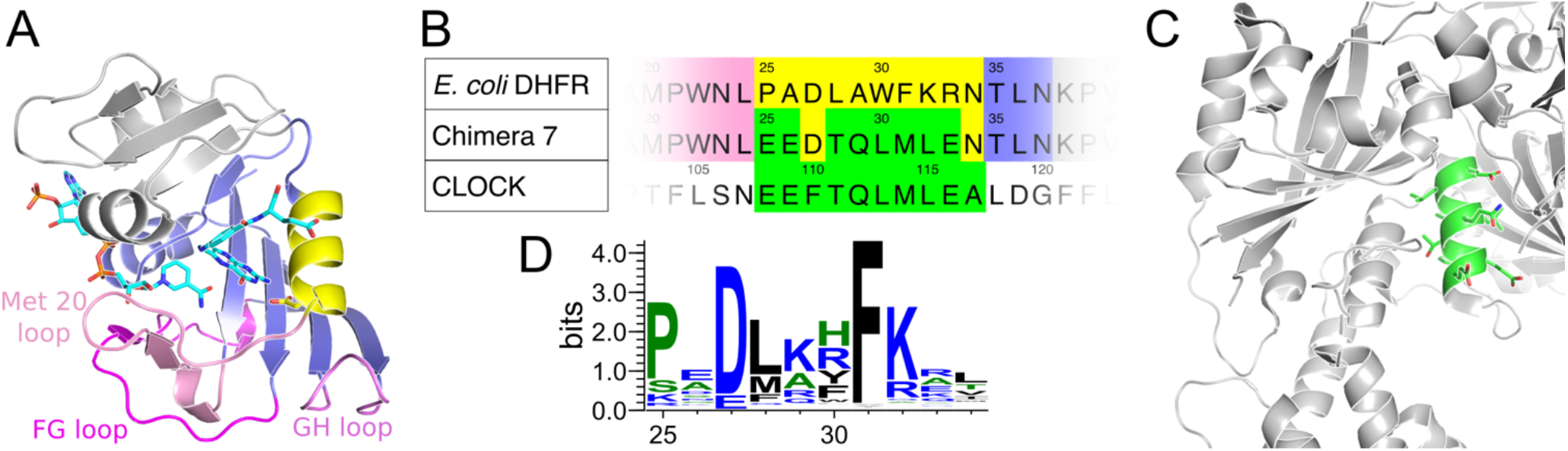
Overview of *E. coli* DHFR and chimera generation. (A) Structure of *E. coli* DHFR (PDB ID 4NX6), with NADP+ and folic acid in teal, adenosine-binding subdomain in gray, and major subdomain in purple and various other colors (Met 20, FG, and GH loops in magenta shades and the α_B_-helix in yellow). The D27 side chain is illustrated using stick rendering. (B) Chimera 7 as an example: alignment of *E. coli* DHFR wildtype, Chimera 7, and helix donor (*M. musculus* CLOCK) sequences. In Chimera 7, the donor helix sequence from CLOCK (green) takes the place of residues 25-34 (yellow) in *E. coli* DHFR, with residues 27 and 34 maintaining their *E. coli* wildtype identities. (C) Structure of CLOCK (PDB ID 4F3L), donor protein for Chimera 7. Residues 108-117, highlighted in green, are substituted into E. coli DHFR to create Chimera 7. (D) Sequence logo for the region of *E. coli* DHFR undergoing replacement, generated by WebLogo (34). Input consists of 19,912 sequences homologous to DHFR, found and aligned by EVcouplings (35).

The chimera generation process is illustrated in Fig. 1B and Fig. 1C using Chimera 7 as an example. The donor sequence, from a helix in *M. musculus* protein CLOCK, replaces nearly all of the *E. coli* DHFR α_B_-helix sequence. We deliberately chose proteins with completely different functions unrelated to the function of DHFR. The chimeric proteins that we studied included non-orthologous helices from other bacteria, *Drosophila*, plants, mammals including humans, as well as transposable elements and mitochondria (see Table S1 in the Supporting Material for all sequences and their source proteins). The catalytically essential residue, D27, was left unmutated for all chimeras. Additionally, in the case of Chimera 7, N34 is also unchanged. For 4 of 20 chimeras, including Chimera 7, some C-terminal positions were left unmutated, to vary the length of the replacement region. Fig. 1D shows a DHFR sequence logo (34) for this 10-residue segment, based on multiple sequence alignment of 19,912 sequences homologous to DHFR found using the EVcouplings web server (35). Several positions (P25, D27, L28, F31, and K32 in *E. coli* DHFR) show a substantial preference for specific residues. The sequence identities of the replacement helical sequences with the original wildtype sequence range from 10% to 50%. In a previous study, we studied six chimeras in which the same residues 25-34 of *E. coli* DHFR were replaced with sequences from six DHFR orthologs (32). We refer to these original six chimeras as the orthologous chimeras and the new chimeras generated in this study as the non-orthologous chimeras.

### Overexpression and purification of DHFR chimeras

See Extended Methods in the Supporting Material for creation of DHFR chimeric constructs. DHFR chimeras fused to C-terminal (6x) His-tag were overexpressed using pFLAG expression vector under isopropyl b-D-1-thiogalactopyranoside (IPTG) inducible T7 promoter described previously (38). Transformed BL21-Gold (DE3) cells were grown in Terrific Broth media (BD) until the cultures reached an optical density of ~0.5, and protein expression was induced overnight at 18°C following the addition of 100μM IPTG. The recombinant proteins were purified from lysates on Ni-NTA columns (Qiagen).

### Steady-state kinetic measurements

DHFR kinetic parameters were measured by progress-curve kinetics as previously described (38). The measurements were carried at 25°C in a 96-well plate format, and the decrease in absorbance at 340 nm due to NADPH oxidation was monitored spectrophotometrically using a Biotek Powerwave HT plate reader. Each assay contained 50 mM MES (2-(N-morpholino)ethanesulfonic acid) pH 7.0, 1mM dithiothreitol, and 150 μM NADPH and purified DHFR chimeras. The reactions were started by the addition of 10/60 μM dihydrofolate (100 μL final volume in the assay).

### Thermal denaturation

Melting temperatures were determined by differential scanning fluorimetry using the fluorescent dye Sypro Orange (Invitrogen). DHFR solutions (5 μM) were prepared in 50 mM phosphate buffer, 1 mM DTT at pH 7.0 in the presence of 100 μM NADPH, and the dye sypro orange was added at 0.5 × final concentration). Thermal denaturation was performed using a Biorad CFX96 Touch with a temperature ramp of 1 °C/min, set between 25 and 90°C, and fluorescence intensity was measured every 0.2°C using the instrument’s FRET channel. The denaturation curves were analyzed using the online software Calfitter v1.3 (39).

### Sequence evolutionary analysis

The probability that a given sequence, {σ}, belongs to an ensemble of aligned, homologous sequences with selected biophysical or functional properties can be modeled using a Boltzmann distribution (40, 41). For instance, direct coupling analysis uses the following model (42, 43):

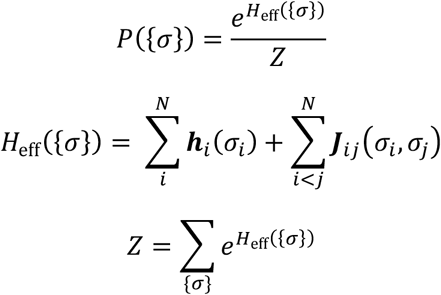

where *N* is the alignment length, ***h**_i_* and ***j**_ij_* are site-specific bias and site-site coupling terms, respectively, and *Z* is the normalizing factor. ***j**_ij_* accounts for epistatic interactions between sites by weighing occurrences of specific residue pairs among the aligned sequences. The parameters are fit using a multiple sequence alignment of homologous sequences. This method has been used to predict the fitness effect of mutations as the difference in the statistical energy (*H*_eff_) between two sequences (44) and is available as the EVcouplings web server and Python package (35). Mutational effects, or Δ*E*, are calculated as Δ*E* = *H*_eff_({σ^ch^}) − *H*_eff_({σ^wt^}). We calculated mutational effects using the epistatic model described above. We also used a non-epistatic model which has a similar form but does not include ***j**_ij_* terms: 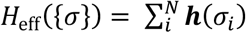. This simpler model reflects sequence conservation at each position (44). In both cases, a more negative Δ*E* indicates a sequence with residues that occur infrequently among DHFR homologs. Δ*E*, therefore, reports on the evolutionary compatibility of replacement fragments with DHFR.

We queried the EVcouplings web server (35) with the *E. coli* DHFR sequence, which yielded 19,912 aligned sequences, and we then used plmc from the Marks Lab (44) to obtain the parameters ***h**_i_* and ***j**_ij_*, calculated over 12,682 effective (non-redundant) sequences. The EVcouplings Python package (35) was used to calculate mutational effects, or Δ*E*. To calculate EVcouplings Δ*E* for random protein sequences, *E. coli* K-12 MG1655 coding sequences (assembly ASM584v2) were obtained from the RefSeq database (45). For all 10-residue subsequences in all non-pseudo gene coding sequences, EVcouplings Δ*E* values were calculated for the substitution of each subsequence for residues 25-34 in *E. coli* DHFR, while preserving position 27 as D.

### Molecular dynamics simulation

We simulated all 27 DHFR variants using replica exchange with solute tempering (REST2) molecular dynamics (MD) simulation. REST2 is an enhanced sampling simulation method in which different replicas use scaled versions of the potential energy (46). Whole-protein REST2 simulations have been previously used to compare DHFR from mesophilic and thermophilic species (47). To avoid inconsistency in folate binding between chimeras, all proteins were simulated in the apo state. For a replica *m*, the potential energy function in a part of the system is scaled by *λ_m_* ≡ *β_m_*/*β*_0_:

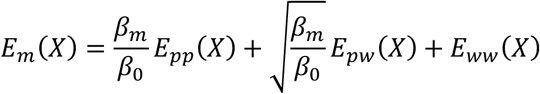

where *w* corresponds to the remainder of the system outside the scaled region of interest, *p*. All replicas are simulated at (inverse) temperature *β*_0_ = 1/*k*_B_T_0_. *β_m_* represents an effective temperature for replica *m* within the scaled region. Replica exchange solute tempering simulations used an effective temperature ladder of 300-550 K and an exchange interval of 500 steps (1 ps). The minimum length of simulation was 300 ns, with simulations of some variants extended to 375 ns to achieve simulation convergence. All replica exchange simulations used 8 replicas, and the acceptance rate between the two lowest rungs was at least 0.15 (acceptance rates varied by DHFR variant). See Extended Methods for details on structure preparation, simulation settings, and simulation analysis.

### Chromosomal replacement of non-orthologous chimeras

Non-orthologous chimeric *E. coli* strains were created by lambda-red recombination (48) according to a previously described procedure (49). The chimeric sequences were introduced by in a pKD13-derived plasmid containing the entire regulatory and coding sequence of the folA gene, flanked by kanR and cmR gene markers and approximately 1 kb homologous region of both upstream and downstream chromosomal genes flanking folA gene. The entire cassette was amplified using primers PCRseq_KefC_for2 and PCRseq_apaH_rev and transformed into BW25113 cells with induced Red helper plasmid pKD46, and cells were recovered in SOC medium containing 0.25x folAmix (adenine 5 ug/mL, inosine 20 ug/mL, thymine 50 ug/mL, methionine 5 ug/mL and glycine 5 ug/mL). Transformants were selected in agar plates containing kanamycin, chloramphenicol, and folAmix. To confirm the correct integration of the desired mutations, folA locus was amplified by PCR (primers Ampl_RRfolA_for, Ampl_RRfolA_rev) and Sanger-sequenced using primer PCRseq_RRfolA_rev. Plasmid pKD46 was removed by plating cells at 37 °C twice in the absence of antibiotic selection. The “wild type strain” referred to in this study represents a BW25113 strain in which the original folA locus was likewise replaced by a replacement cassette containing the sequence of wild type DHFR flanked by kanR and cmR antibiotic resistance markers. Three colonies of each chimeric or wild type *E. coli* strain were picked from agar plates and grown in M9 minimal media supplemented with 0.25x folAmix until saturation. Glycerol was added (10% final volume) to the grown cultures and various aliquots were stored at −80°C for subsequent studies.

### Growth rate

Overnight cultures were prepared by inoculating 200 μL of M9 minimal media. supplemented with 0.8 g/L glucose, 1mM MgSO_4_, 0.1mM CaCl_2_, 50 μg/mL kanamycin, 34 μg/mL chloramphenicol and 0.25x folAmix, with 5 uL of thawed −80 °C glycerol stocks. The next morning, the cultures were diluted into fresh M9 minimal media (composition as above) to an optical density (OD) of approximately 0.01. After a period of 6-8h growth, the cultures were washed with M9 minimal medial without folAmix by 3 cycles of centrifugation/resuspending the cell pellets 3 times. After the final resuspension, the OD of the cultures was normalized to 0.0015. A 96-well microplate containing 100 uL of M9 minimal media, supplemented with 0.8g/L glucose, 1mM MgSO_4_, 0.1mM CaCl_2_, 50 μg/mL kanamycin, 34 μg/mL chloramphenicol, and different concentrations of folAmix, was inoculated with 50 uL/well of the normalized cultures. The growth measurements were performed at 37°C with a Tecan Infinite M200 Pro microplate reader.

### Intracellular protein abundance

The intracellular abundance of chimeric DHFR was determined by Western blotting. Chimeric strains were grown at 37°C in M9 minimal media supplemented with 2 g/L glucose, 50 μg/mL kanamycin, 34 μg/mL chloramphenicol, and 0.25x folAmix. When the optical density (OD) was approximately 0.3 the cultures (5mL) were centrifuged and the supernatant discarded. Based on the OD of the cultures before centrifugation, the pellets were resuspended to an OD of 11 in lysis buffer consisting of 100mM HEPES pH 7.3, 1x Popculture reagent (Millipore), 10x Complete protease inhibitor cocktail (Sigma), 1 mg/mL lysozyme, 1U/mL Benzonase (Sigma) and 1mM DTT. The tubes were sonicated in a water bath for 20-30 min until complete lysis and the insoluble material was removed by centrifugation at 20000xg for 10 min. 30 μL of the cleared supernatants were mixed with 15 μL SDS-PAGE loading buffer (3X) containing β-mercaptoethanol. The samples were incubated for 20 min at 37°C and loaded on an SDS-PAGE gel together with colored MW markers. After the separation, the gel was cut in half and the low-MW part (below 37kDa) was transferred to a nitrocellulose membrane. The DHFR-specific band was identified using rabbit polyclonal anti-DHFR antibodies (1/20000 dilution) followed by alkaline phosphatase-linked anti-rabbit secondary antibodies (1/10000 dilution). The band was revealed with Novex™ AP Chromogenic Substrate (BCIP⁄ NBT) (Thermofisher Scientific). The other half of the SDS-PAGE gel with the high-MW bands was stained with Bio-Safe™ Coomassie Stain (Biorad). The intensity of Western-blot bands was quantified by the ImageJ software (50), and was normalized by the total protein content, which was estimated by the total intensity of each corresponding lane in the Coomassie-stained high-MW half of the SDS-PAGE gel.

### DHFR fitness model

A fitness model for DHFR was described previously (38). The flux through DHFR enzymatic reaction was estimated for each chimeric strain using the equation,

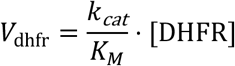

where [DHFR] is the intracellular concentration determined for the chimeric protein. Flux values were normalized with respect to that of the wild type strain: *V*_norm_ = *V*_dhfr_ / *V*_dhfr,WT_. The model posits growth rate, *y*, is a function of flux, with the following Michaelis-Menten-like form:

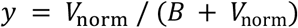

where *B* = 0.001. To estimate the growth rate at different levels of flux, growth rates for the wild type strain were measured at different concentrations of trimethoprim (38), and the calculated flux was corrected by a factor, 1 / (1 + α٠[TMP]_medium_ / *K_i_*), where [TMP]_medium_ is the concentration of trimethoprim in the growth medium, α is the ratio between the intracellular trimethoprim concentration and that of the growth medium, and *K*_i_ is the trimethoprim inhibition constant (0.94 nM for wild type DHFR). The parameter α must be determined independently for different strains since cell permeability and overall sensitivity to trimethoprim are expected to vary. For the strain BW25113 used in this work, α = 0.4.

## Results

### Non-orthologous DHFR chimeras have reduced catalytic activity

Twenty chimeric, non-orthologous DHFR variants were overexpressed in *E. coli*, purified to homogeneity, and characterized biophysically and biochemically. The melting temperature (*T_m_*) of each chimera was measured by following the changes in intrinsic tryptophan fluorescence upon thermal denaturation. Except for Chimera 4, the thermal stabilities of the non-orthologous chimeras are reduced (Fig. 2). For 14 of 20 non-orthologous chimeras, *T_m_* is at least 10 °C lower than that of the wild type (WT), and the majority have *T_m_* values lower than those of the orthologous chimeras. Size exclusion chromatography revealed that all tested chimeras elute as monomers, except for Chimera 5, which elutes as a mixture of monomer and dimer (Fig. S1 in the Supporting Material). The relatively high *T*_m_ (54.4 °C) that was measured for Chimera 5 is likely associated with additional stabilization conferred by dimerization.

**Fig. 2:**
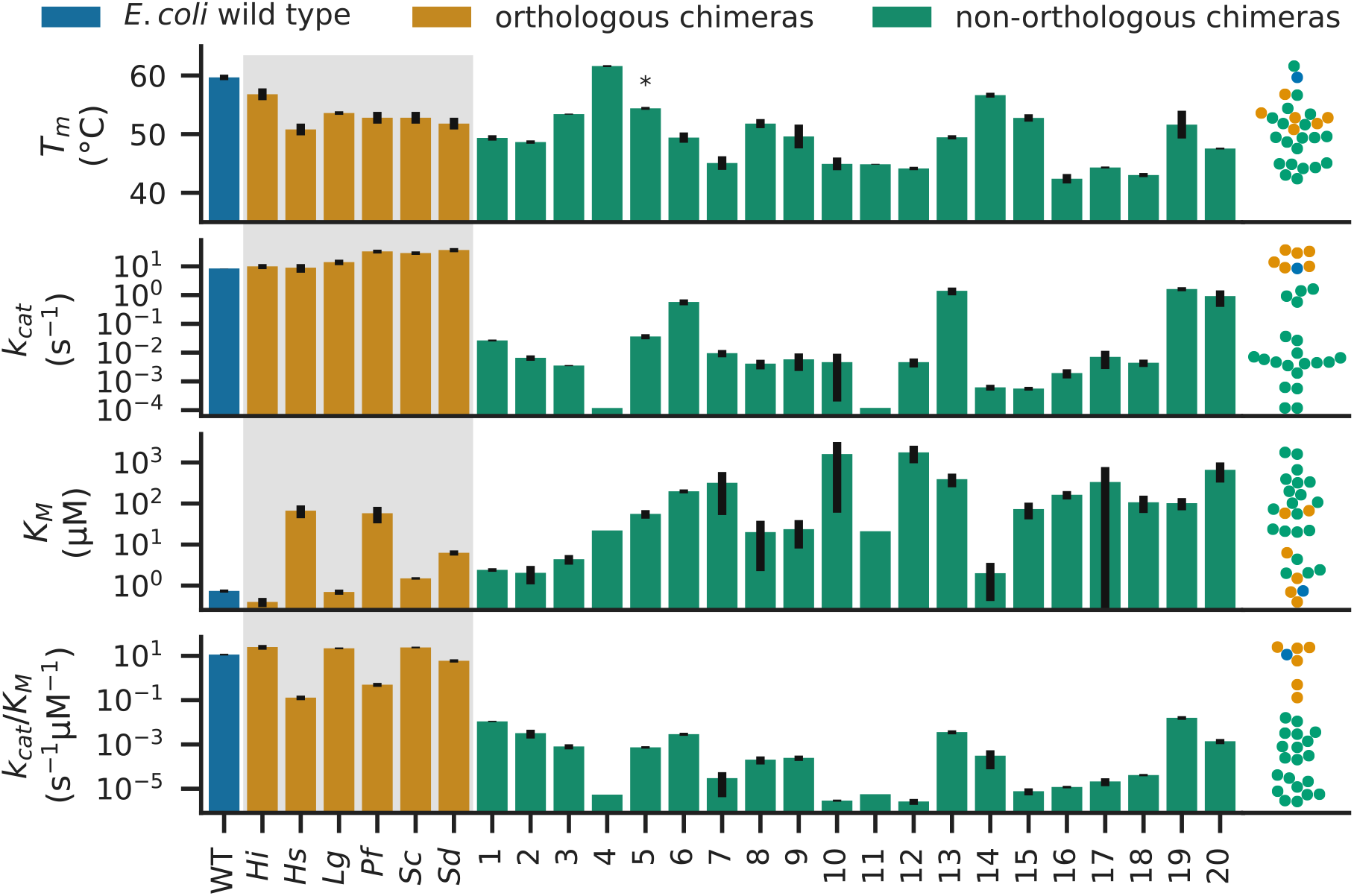
Stabilities and catalytic activities of non-orthologous chimeras of DHFR variants are generally decreased. Error bars indicate the standard error of the mean. Right-hand side swarm plots show distributions of corresponding properties (values are plotted against the vertical axis with horizontal offsets to avoid overlap). Orthologous chimeras (orange bars, gray background) were characterized in previous work (32). *: Size exclusion chromatography found that Chimera 5 is partly dimeric (Fig. S1).

Substitution with non-orthologous helical sequences compromises the catalytic function of the protein. The catalytic rates (*k_cat_*) and catalytic efficiencies (*k_cat_*/*K_M_*) of non-orthologous chimeras are all lower than those of wildtype DHFR and the orthologous chimeras (Fig. 2). *k_cat_*/*K_M_* values are at least three orders of magnitude lower than that of wildtype DHFR. Substrate affinity (*K_M_*) is also reduced (higher values) for non-orthologous chimeras. One possible reason for the differences among the chimeras could be that the replacement segments differ in their helical propensities. We found, however, no correlations between measured *in vitro* properties and the Agadir score (Fig. S2 in the Supporting Material), a computational prediction for the helical propensity of the sequences in isolation (51–55). This suggests that a mechanism that accounts for interactions with the rest of the protein is more likely to explain the observed differences.

### Helix residues are under evolutionary selection for catalytic activity

The striking difference in catalytic activity (several orders of magnitude) between orthologous and non-orthologous chimeras suggests that all or some residues in the α_B_-helix were evolutionarily selected to maintain the catalytic activity of the enzyme. To this end, we assessed how divergent our replacement sequences are from DHFR using the evolutionary analysis tool, EVcouplings (35). We calculate a statistical Δ*E* that describes the probability of a sequence belonging to a family of homologous sequences, with more negative Δ*E* indicating lower probability.

EVcouplings non-epistatic model Δ*E* has strong correlations with catalytic parameters *k_cat_* and *k_cat_*/*K_M_*. Scatterplots between the non-epistatic Δ*E* and experimental properties are shown in Fig. 3. While Spearman correlations between Δ*E* and *T_m_* (*ρ* = 0.38, *P* = 0.049, Fig. 3A) and between Δ*E* and *K_M_* are weak (*ρ* = −0.28, *P* = 0.16, Fig. 3B), there are strong correlations between Δ*E* and *k_cat_* (*ρ* = 0.84, *P* < 0.001, Fig. 3C) and between Δ*E* and *k_cat_*/*K_M_* (*ρ* = 0.74, *P* < 0.001, Fig. 3D). Similar but slightly weaker correlations are obtained using Δ*E* according to the epistatic model (Fig. S3 in the Supporting Material). These correlations relate evolutionary data to experimental protein properties to indicate that the residues of the α_B_-helix have evolved under selection pressure to maintain a high enzyme catalytic rate and efficiency. The correlations between catalytic activity and Δ*E* are stronger than the correlations calculated using sequence identity (*k_cat_* and sequence identity: *ρ* = 0.63, *P* < 0.001) or sequence similarity (*k_cat_* and sequence similarity: *ρ* = 0.63, *P* < 0.001) (Table S2 and Fig. S4 in the Supporting Material). Sequence similarity was calculated according to Grantham’s distance between amino acids, which is a model based on residue properties (56).

**Fig. 3:**
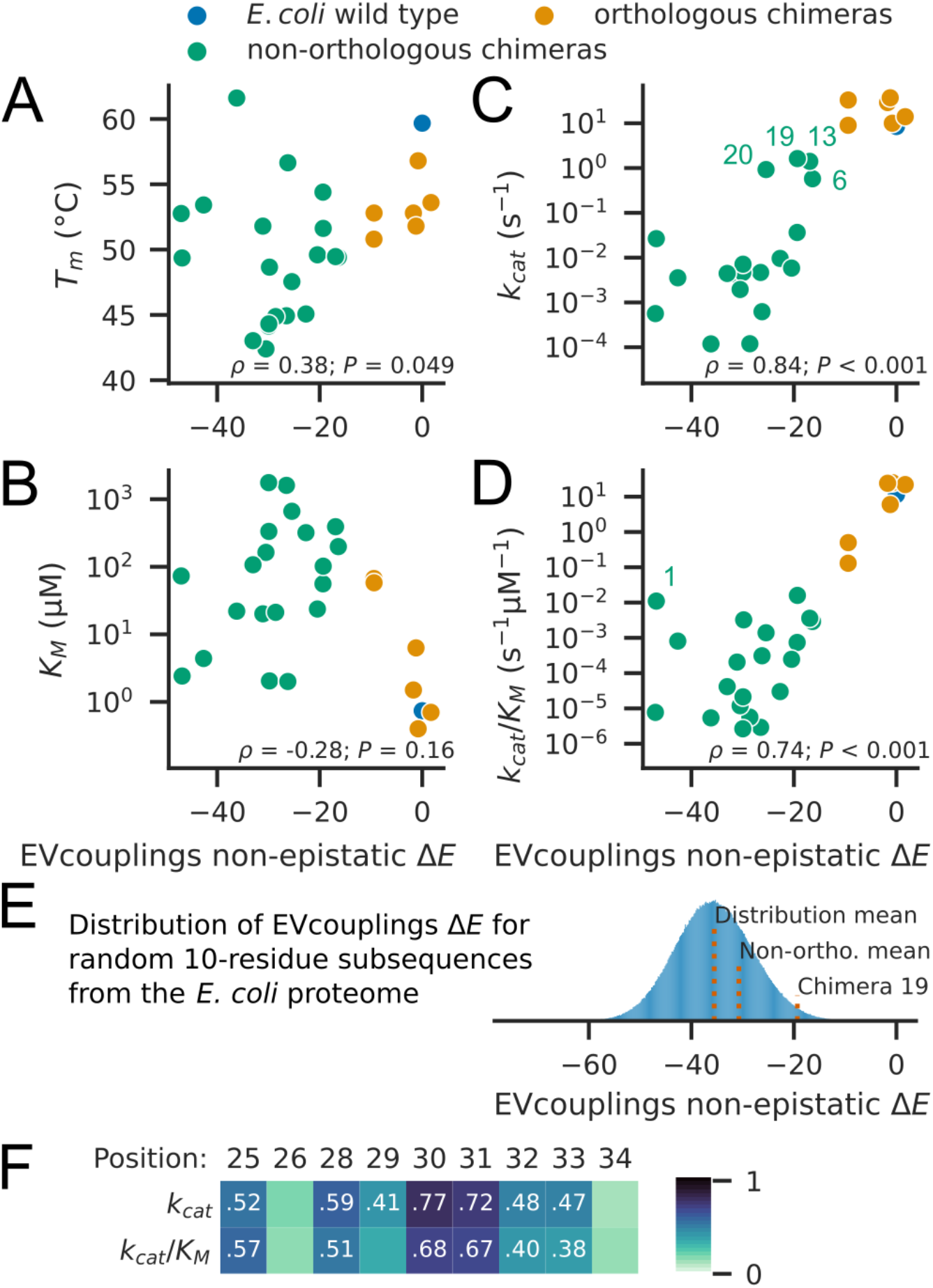
EVcouplings non-epistatic model Δ*E* correlates with catalytic turnover and efficiency of DHFR chimeras. Non-epistatic Δ*E* is compared with (A) *T_m_*, (B) *K_M_*, (C) *k_cat_*, with Chimeras 6, 13, 19, and 20 labeled, and (D) *k_cat_*/*K_M_*, with Chimera 1 labeled. Spearman rank correlation coefficients and *P* values are given in the lower portion of each plot. (E) Distribution of non-epistatic Δ*E* from substituting α_B_-helix residues 25-34 in *E. coli* DHFR with all 10-residue subsequences from all *E. coli* coding sequences while retaining D27 (n = 1282235). The mean of the distribution, the mean Δ*E* from the 16 non-orthologous chimeras (Non-ortho.) in which residues 25-34 were substituted (excludes Chimeras 6, 7, 8, and 13), and Δ*E* for Chimera 19 are indicated. (F) Heatmap of Spearman rank correlation coefficients between catalytic parameters, *k_cat_* and *k_cat_*/*K_M_*, and non-epistatic Δ*E* values of point mutations at each position. Correlation coefficients are printed for statistically significant (*P* < 0.05) combinations.

Examining individual chimeras, we find that four chimeras have at least one order of magnitude higher *k_cat_* than those of all other non-orthologous chimeras: Chimeras 6, 13, 19, and 20 (Fig. 3C). Chimeras 6 and 13 have higher wildtype sequence identities (40% and 50%, respectively), as a result of shorter helix replacement lengths (25-32 and 25-30, respectively), which likely explain their higher *k_cat_* values and less negative Δ*E* values. By contrast, Chimeras 19 and 20 both have only 20% identity. Chimera 19 notably has the highest *k_cat_* and *k_cat_*/*K_M_* of all non-orthologous chimeras. The Chimera 19 sequence matches the wildtype DHFR sequence at position 30 (W30); this, alongside the overall compatibility of other residues, likely explains its high activity. Additionally, in considering *k_cat_*/*K_M_* (Fig. 3D), one outlier stands out: Chimera 1, which has the second-lowest non-epistatic model Δ*E* and the second-highest *k_cat_*/*K_M_* among non-orthologous chimeras. The α_B_-helix sequence for chimera 1, VQDDTEELKS^25-34^ (10% identity), switches residue hydrophobicity at positions 28, 30, 31, and 32, all sites with substantial sequence conservation (Fig. 1D). The relatively high activity of this variant is therefore surprising and is further explored in the next section.

To contextualize our helix sequences and to gauge the rarity of sequences like Chimera 19, we calculated non-epistatic Δ*E* values for substitution of α_B_-helix residues 25-34 with all valid 10-residue subsequences (n = 1282235) from protein-coding sequences in *E. coli*, while keeping position 27 as D. We find that Chimera 19-like sequences are rare: 1.80 % of the sequences have a higher Δ*E* than that of Chimera 19 (Fig. 3E). We compared these protein subsequences to the 16 non-orthologous helical sequences in which residues 25-34 were substituted. The non-orthologous chimeras have a slightly less negative mean Δ*E* than that of random protein subsequences (−30.7 versus −35.7). The difference between the distributions was statistically significant at the *P* = 0.05 level by Mann-Whitney *U* test (*P* = 0.0058) and Kolmogorov-Smirnov test (*P* = 0.0027). The data suggest that α-helical sequences have slightly better compatibility upon substitution than random protein segments.

Finally, we examined the contributions to Δ*E* from each position in the α_B_-helix. We calculated the correlation of Δ*E* for point mutations to the chimeric DHFR residues at each α_B_-helix position with *k_cat_* and *k_cat_*/*K_M_* of the chimeras. For all sites except positions 26 and 34, there are statistically significant correlations between point mutation effects and at least one of *k_cat_* and *k_cat_*/*K_M_* (Fig. 3F). Mutational effects at positions 30 and 31 have the strongest correlations with *k_cat_* and *k_cat_*/*K_M_*. The residue identities of positions 30 and 31 appear most important in determining the catalytic activity of helix-swapped DHFR chimeras. These two sites are inward-facing and hydrophobic in the wildtype protein (W30 and F31, respectively), with F31 forming part of the substrate-binding pocket. Notably, Chimera 19 retains W30, and all orthologous chimeras retain F31. Overall, EVcouplings analysis shows that the residues of the non-orthologous replacement sequences are less likely to appear among sequences homologous to DHFR, and our *in vitro* measurements of catalytic activity indicate that such residues were selected against in evolution due to deleterious effects on catalysis. We next try to understand the low catalytic activity of these DHFR variants on a structural level.

### Simulations find chimeras have destabilized α_B_-helices

To investigate the structures of the DHFR chimeras and understand the physical reason behind their decreased catalytic activity, we turned to REST2 MD simulation. Our analyses use simulation data from the last 100 ns of simulation. Fig. S5 in the Supporting Material demonstrates that the root mean square deviation (RMSD) of the Cα atoms of the α_B_-helix reaches steady state over at least the last 100 ns of simulations. The RMSD of the α_B_-helix is measured after alignment of the major subdomain core residues (the β-sheet), using the DHFR crystal structure as the reference. This enables detection of conformational differences in which the entire α_B_-helix is displaced relative to the core of the protein.

The non-orthologous chimeras generally show greater deviation from the *E. coli* DHFR crystal structure reference. Fig. 4A compares the mean α_B_-helix RMSDs of wildtype DHFR and orthologous chimeras and non-orthologous chimeras. All six orthologous chimeras maintain the α_B_-helix in the wildtype-like conformation despite three of the six orthologous chimeras having only 20% α_B_-helix sequence identity with *E. coli* DHFR, demonstrating evolutionary structural conservation. Only the *P. falciparum* (*Pf*) orthologous chimera shows a slightly higher helix RMSD (1.73 Å). By contrast, just 4 of the 20 non-orthologous chimeras (Chimeras 2, 13, 18, and 19) have mean helix RMSDs approaching those of wildtype DHFR and the orthologous chimeras.

**Fig. 4:**
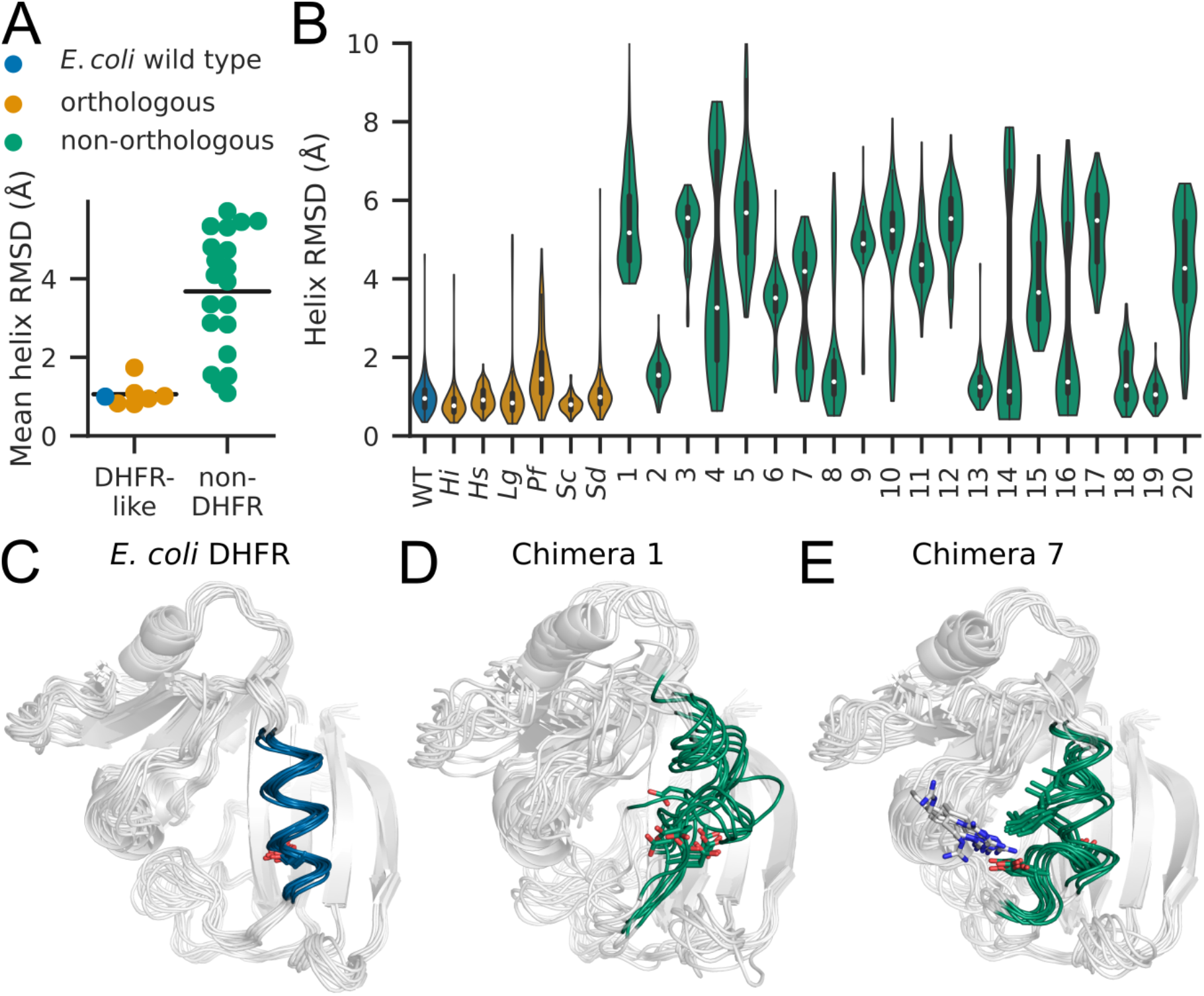
Replica exchange MD simulations of DHFR variants show non-orthologous chimeras deviate from *E. coli* wildtype conformations. (A) Mean helix RMSDs of *E. coli* wildtype DHFR and orthologous chimeras (DHFR-like) are generally lower than those of non-orthologous chimeras (non-DHFR). Horizontal lines indicate the mean value for each group. (B) Violin plots of helix RMSD show distributions of helix RMSD for each chimera. Inner white marker denotes median value, and ends of thicker vertical bars denote 25^th^ and 75^th^ percentiles. (C, D, and E) Superimposed structures of *E. coli* DHFR, Chimera 1, and Chimera 7 taken from the last 100 ns of each simulation, 10 samples per image, evenly separated in time. Helix backbones (25–35) are colored and depicted in loop representation. The D27 sidechains of all three variants as well as the R52 and L32 sidechains of Chimera 7 are illustrated using stick representation. Samples have been aligned using PyMOL (57).

The distributions of helix RMSDs of the DHFR variants (Fig. 4B) suggest that the non-orthologous chimeras can be divided into two groups, those with predominantly helical, wildtype-like α_B_-helix conformations and those exhibiting non-wildtype conformations. The median helix RMSD (Fig. 4B) is one differentiator; seven chimeras have values below 2 Å. The α_B_-helices in these chimeras are fully helical in the majority of simulation frames: the median percent helicities of these chimeras are all 100% (Fig. S6 in the Supporting Material), as calculated by DSSP (58, 59). Compared to the orthologous chimeras, the α_B_-helices in these wildtype-like non-orthologous chimeras can be subtly different from the wildtype structure or exhibit more disordered conformations, which both result in higher mean helix RMSDs. Nevertheless, Chimera 19, which has the highest *k_cat_* and *k_cat_*/*K_M_* of all non-orthologous chimeras, is particularly wildtype-like (1.09 Å mean helix RMSD), demonstrating one case in which structure and function can be maintained after a helix swap with an evolutionarily unrelated sequence.

The second category of non-orthologous chimeras shows greater disorder in the α_B_-helix. Two examples from this second group of non-orthologous chimeras, Chimeras 1 and 7, are compared to wildtype DHFR in Fig. 4C, Fig. 4D, and Fig. 4E. Each image shows 10 samples from the last 100 ns of simulation. While wildtype DHFR maintains the crystallographic helix conformation, with D27 in its ligand-binding orientation (Fig. 4C), Chimeras 1 and 7 both show ensembles of more variable helix conformations, and the three loops of the major subdomain are also visibly more disordered (Fig. 4D and Fig. 4E, respectively). Chimera 1 maintains partial helicity in just the C-terminal half of the α_B_-helix, and the D27 sidechain faces outward (Fig. 4D). Since Chimera 1 has relatively high catalytic activity (second highest *k_cat_*/*K_M_* among non-orthologous chimeras), it is remarkable that its α_B_-helix conformations are among the least wildtype-like (second highest mean helix RMSD). These observations are unlikely to be due to simulation inaccuracy given that the Chimera 1 α_B_-helix sequence contains several polar residues (Table S1). Secondly, Chimera 7 shows both wildtype-like and non-wildtype α_B_-helix conformations (Fig. 4E). The median helix RMSD of 4.2 Å for Chimera 7 (Fig. 4B) indicates, however, that the non-wildtype conformations are more populated than the wildtype-like conformations. The non-wildtype conformation is characterized by occlusion of the active site as a result of L32 occupying the active site pocket and D27 interacting with R52 (Fig. 4E). Since loss of the wildtype α_B_-helix conformation can block the binding site and disrupts the positioning of the catalytically essential D27 (Fig. S6), the non-wildtype helix conformations in non-orthologous chimeras likely explains a large part of the loss of catalytic activity. We next examine the correlation of these simulation measurements with experimental properties.

### Simulation measurements correlate most strongly with thermal stability and K_*M*_

The different conformations of the DHFR variants prompted an investigation of the correlation between simulation measurements and *in vitro* properties of our variants. In simulations, non-helical conformations in the replacement segment are often characterized by the burial of hydrophobic sidechains into the substrate binding pocket or the hydrophobic surface of the major subdomain β-sheet, as was seen for Chimera 7 (Fig 4E). We thus examined differences in hydrophobic solvent accessible surface area (SASA) among the DHFR variants. This quantity was measured by summing up the SASA of nonpolar atoms belonging to the α_B_-helix. This measurement is done in the context of the full protein structure; buried nonpolar helix atoms do not have any contribution to hydrophobic SASA (see Extended Methods). Helix hydrophobic SASA has moderately high correlations with both *T_m_* (*ρ* = 0.78, *P* < 0.001, Fig. 5A) and *K_M_* (*ρ* = −0.65, *P* < 0.001, Fig. 5B). That is, DHFR variants with greater solvent-exposed hydrophobic surface areas on helix residues have better thermal stabilities and substrate binding affinities. This is a trend that both orthologous and non-orthologous chimeras follow, and correlations remain moderately high and statistically significant for both *in vitro* quantities when only non-orthologous chimeras are considered (helix hydrophobic SASA and *T_m_*: *ρ* = 0.67, *P* = 0.0013; helix hydrophobic SASA and *K_M_*: *ρ* = −0.63, *P* = 0.0027). *T_m_* and *K_M_* are correlated properties (*ρ* = −0.68, *P* < 0.001), with more destabilized DHFR variants also having higher *K_M_* values (Fig. S7 in the Supporting Material). Decreased thermal stability likely reduces substrate affinity by making the optimal substrate-binding conformation less probable. But it is unintuitive that DHFR variants with more solvent-exposed nonpolar surface area would be associated with greater thermal stability. We note that helix hydrophobic SASA is not an indicator of wildtype-like α_B_-helix conformations, since the correlation between helix hydrophobic SASA and helix RMSD is weak (*ρ* = −0.21, *P* = 0.28, Fig. S8 in the Supporting Material).

**Fig. 5:**
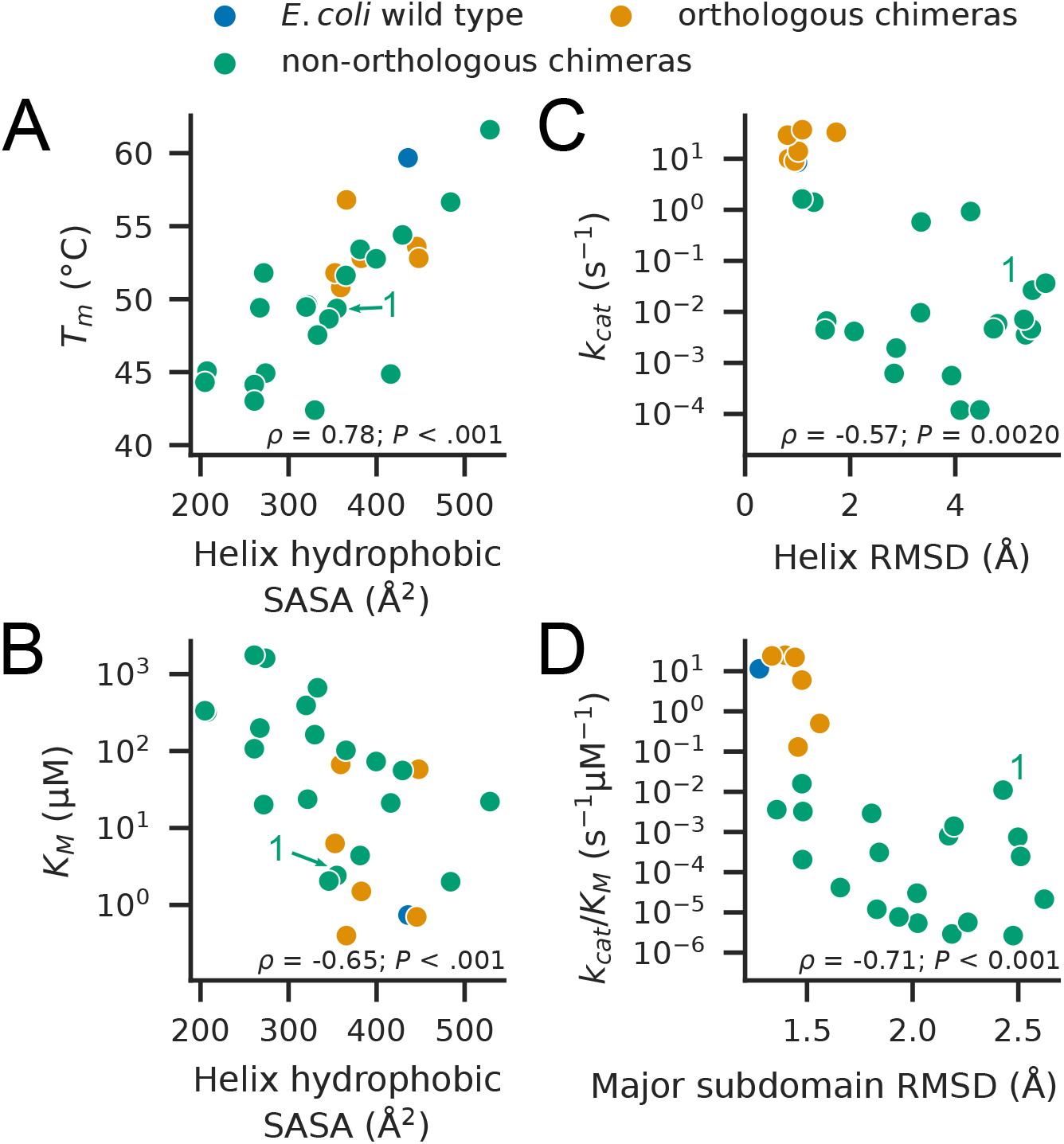
Measures of hydrophobic solvent accessible surface area (SASA) correlate with *T_m_* and *K_M_*, and RMSD measurements correlate with *k_cat_* and *k_cat_*/*K_M_*. Scatterplots between (A) helix hydrophobic SASA and *T_m_*, (B) helix hydrophobic SASA and *K_M_*, (C) α_B_-helix RMSD and *k_cat_*, and (D) major subdomain RMSD and *k_cat_*/*K_M_*. Simulation quantities are mean values. Chimera 1 is labeled in each plot. Spearman rank correlation coefficients and *P* values are given in the lower portion of each plot. When only non-orthologous chimeras are considered, the correlations with helix hydrophobic SASA remain moderately high and statistically significant: (A) *ρ* = 0.67, *P* = 0.0013, (B) *ρ* = −0.63, *P* = 0.0027, (C) *ρ* = −0.08, *P* = 0.75, (D) *ρ* = −0.36, *P* = 0.12.

We also examined simulation order parameters for different parts of the protein. Differences in disorder among DHFR variants occur in the major subdomain, while the adenosine-binding subdomain remains stable (Fig. S6). The adenosine-binding subdomain is more distant from the α_B_-helix and exhibits greater stability in native state proteolysis experiments (60). We thus focused our analysis on the various regions of the major subdomain. We find statistically significant correlations between mean RMSD measurements of different regions of the DHFR major subdomain and the experimentally determined *in vitro* properties of *T_m_*, *k_cat_*, *K_M,_* and *k_cat_*/*K_M_* (Fig. S9 in the Supporting Material). The RMSD measurements of different regions are also substantially correlated with each other (Fig. S8), reflecting the cooperative nature of protein stability. Fig. 5C and Fig. 5D show the strongest correlations seen in Fig. S9 between the RMSD measurements and *k_cat_* and between the RMSD measurements and *k_cat_*/*K_M_*, respectively. The correlation between helix RMSD and *k_cat_* is moderate (*ρ* = −0.57, *P* = 0.002, Fig. 5C), while the correlation between RMSD of the entire major subdomain and *k_cat_*/*K_M_* is moderately high (*ρ* = −0.71, *P* < 0.001, Fig. 5D). These trends are driven by the difference between orthologous and non-orthologous chimeras, however. Among non-orthologous chimeras, there is no correlation between helix RMSD and *k_cat_* (*ρ* = −0.08, *P* = 0.78), and the correlation between major subdomain RMSD and *k_cat_*/*K_M_* is weak and no longer statistically significant (*ρ* = −0.36, *P* = 0.12). These trends show that while maintenance of the wildtype DHFR conformation is associated with high *k_cat_* and *k_cat_*/*K_M_*, the degree of deviation from the wildtype conformation weakly predicts these catalytic parameters.

The previously discussed Chimera 1 is an outlier with respect to RMSD and *k_cat_*/*K_M_* (Fig. 5D). Chimera 1 also has an unusually low *K_M_* (3^rd^ lowest among non-orthologous chimeras), given its overall deviation from the wildtype conformation (Fig. S9). The conformations of Chimera 1 (Fig. 4D) show that the α_B_-helix conformations in Chimera 1 are not wildtype-like but do leave the active site relatively open. By contrast, many of the other disordered-helix non-orthologous chimeras have conformations that block the DHF binding site. To quantify this, we measured DHF binding site occlusion by overlaying the folate ligand from the *E. coli* DHFR crystal structure into simulation frames after structural alignment using the major subdomain core residues. Occlusion was calculated as the proportion of ligand volume that is occupied by protein atoms (see Extended Methods). Chimera 1 exhibits only moderate occlusion, comparable to chimeras with high helicity and wildtype-like α_B_-helix conformations (Fig. 6A). The wildtype DHFR and Chimera 8 are also notable for low binding site occlusion.

**Fig. 6:**
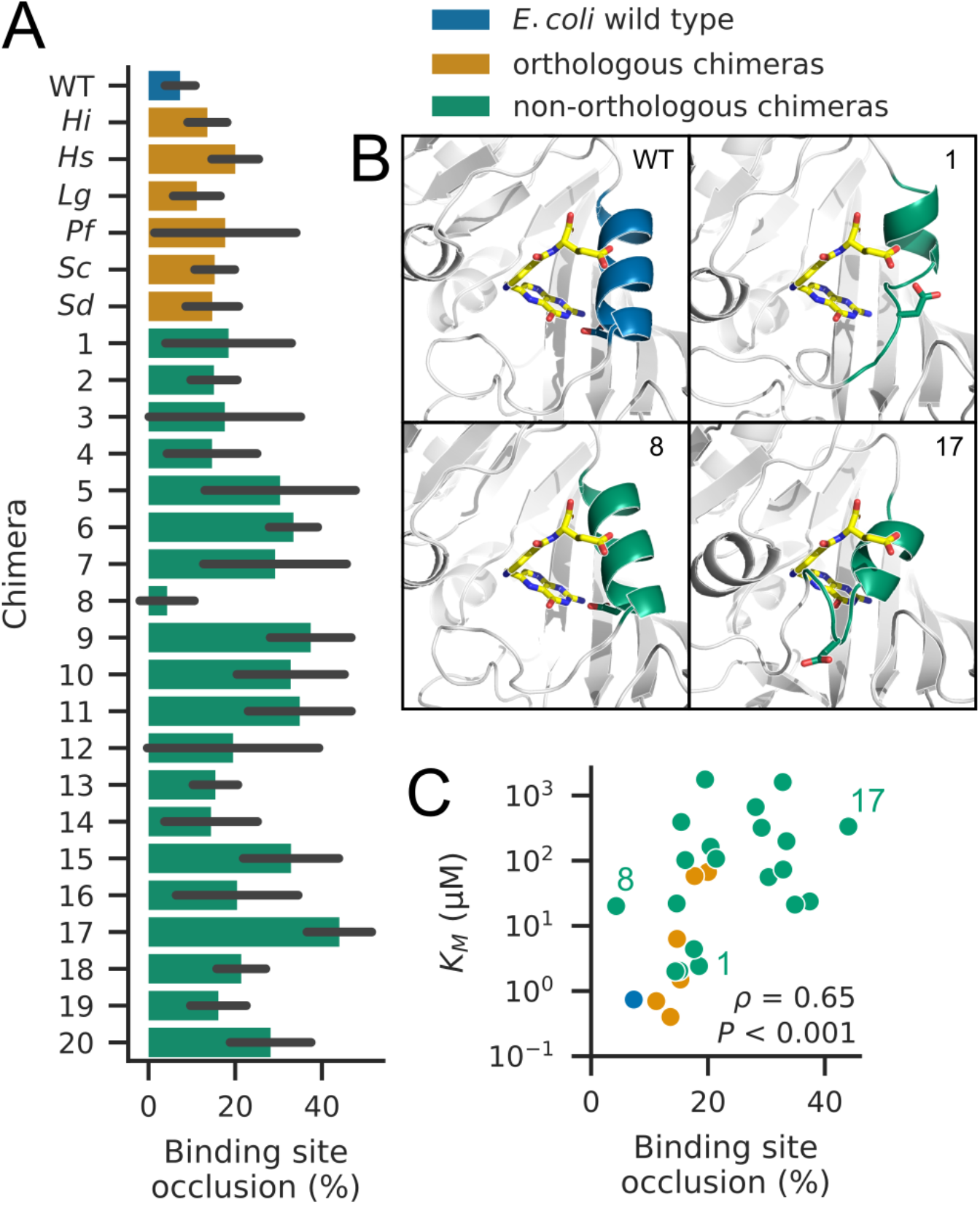
DHFR variants with greater DHF binding site occlusion have higher *K_M_* values. Binding site occlusion was measured by calculating the amount of volume available to the DHF substrate. (A) Average binding site occlusion for each DHFR variant. Error bars represent standard deviations. (B) Representative structures from simulations for *E. coli* wildtype DHFR and Chimeras 1, 8, and 17 are illustrated. Conformations adopted by Chimeras 1 and 8 do not block the substrate binding site, whereas Chimera 17 conformations occupy the substrate binding site. The yellow folate ligand is overlaid from alignment to the DHFR crystal structure using the core residues of the major subdomain, and the α_B_-helix (residues 25-35) is colored. (C) Scatterplot of binding site occlusion versus *K_M_*. Chimeras 1, 8, and 17 are labeled. Spearman rank correlation coefficient and *P* value are given in the lower right of the plot. When only non-orthologous chimeras are considered, correlation is weaker (*ρ* = 0.42, *P* = 0.069).

Fig. 6B highlights representative structures from simulations in which the folate ligand from the DHFR crystal structure is superimposed on the simulation frames for wildtype DHFR and Chimeras 1, 8, and 17. Apo state wildtype DHFR maintains the crystallographic conformation, with D27 well-aligned with the superimposed folate ligand. For Chimera 1, the α_B_-helix residues adopt a partly helical, open conformation, leaving the binding site free. Chimera 8 is also illustrated because it has exceptionally low binding site occlusion, which can be explained by an α_B_-helix conformation that is helical but tilted away compared to wildtype DHFR. By contrast, Chimera 17 exhibits the highest binding site occlusion and typifies how most non-helical conformations block the DHF binding site. The residues of Chimera 17 overlap with the atoms of the superimposed folate ligand. We tested whether binding site occlusion correlates with substrate affinity, *K_M_*, and indeed, we find our measure of binding site occlusion has a moderately high correlation with *K_M_* (*ρ* = 0.65, *P* < 0.001, Fig. 6C), although the correlation is weaker when only non-orthologous chimeras are considered (*ρ* = 0.42, *P* = 0.069). Thus, by maintaining an open DHF binding site, Chimera 1 differs from other non-orthologous chimeras that have disordered α_B_-helices.

### Low catalytic efficiency and decreased intracellular abundance strongly impact the fitness of *E. coli* strains expressing non-orthologous chimeras

To determine the extent of modularity in α-helices and its relevance to evolution, it is crucial to evaluate how changes in protein biophysical properties caused by the swapping of structural segments can affect the fitness of an organism. To investigate this, we carried out the chromosomal replacement of the wildtype DHFR-coding gene in *E. coli* (*folA*) with the sequences of the five non-orthologous chimeras with the highest *k_cat_*/*K_M_* (Chimeras 1, 2, 6, 13, and 19). Anticipating that the low catalytic function of these proteins would make the strains non-viable, all gene editing steps were made in the presence of folAmix, a metabolic supplement that rescues cells from DHFR deficiency (49). Upon successful *folA* gene replacement with Chimeras 1, 6, 13, and 19 (but not Chimera 2), we tested for the ability of these strains to grow in the absence of folAmix supplement. Under such conditions, only Chimeras 13 and 19 are viable, though their growth rates are lower than that of the wild type (Fig. 7A).

**Fig. 7:**
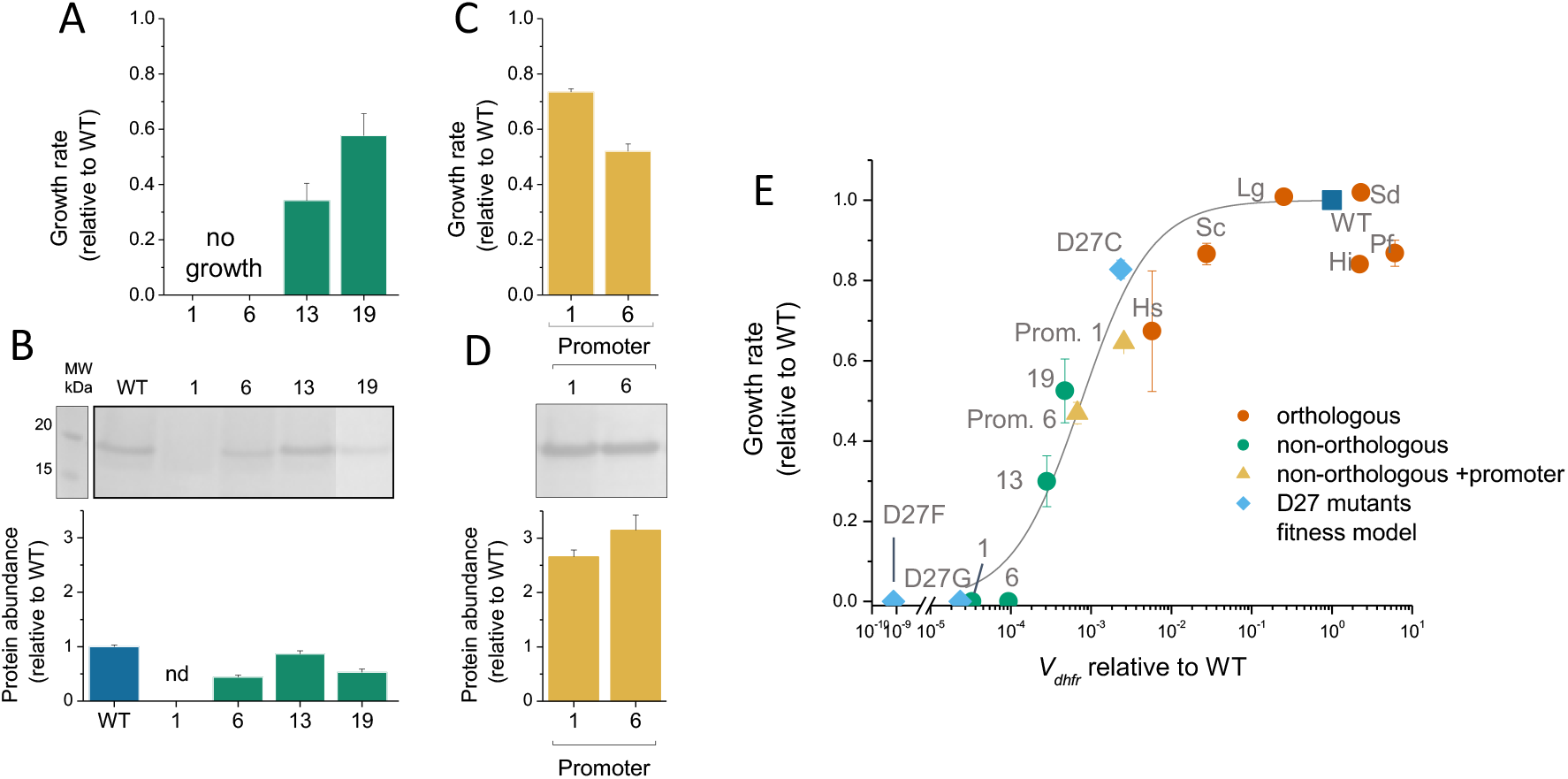
The fitness of non-orthologous chimeric DHFR strains is markedly reduced due to a combination of decreased catalytic activity and low intracellular protein abundance. (A) Chromosomal replacement of the gene *folA* with non-orthologous chimeric sequences produces strains that are non-viable or that have decreased fitness. No detectable growth was observed for strains with Chimeras 1 and 6 after 48 h. The growth rates were normalized with respect to that of the strain with wildtype DHFR, and the represented values are the mean ± standard deviation of three replicates. (B) The intracellular abundance of DHFR is decreased in chimeric strains. The amount of DHFR protein present in the soluble fraction of lysates of chimeric strains was determined by Western blot using anti-*E. coli* DHFR polyclonal antibodies (top panel and Fig. S10). The represented values are the mean ± standard deviation of three replicates and were normalized with respect to the levels of wild type DHFR. (C), (D) A mutation (C-35T) in the promoter region of the *folA* gene that increases the levels of DHFR expression rescues the growth of strains with Chimeras 1 and 6. (E) The fitness determined for chimeric strains agrees with the DHFR fitness model based on the protein biophysical properties (38). *V*_dhfr_, the normalized flux through DHFR enzymatic reaction was estimated for each chimeric strain using the equation *V*_dhfr_ = *k_cat_*/*K_M_*٠[DHFR], where [DHFR] is the intracellular concentration DHFR determined for each chimeric strain. The solid line represents the best fit of the data with the equation *y* = *V*_norm_ / (*B* + *V*_norm_) where *V*_norm_ = *V*_dhfr_ / *V*_dhfr,WT_ and *B* = 0.001 (38).

The observation that Chimera 1 could not grow, despite having higher *k_cat_*/*K_M_* than that of Chimera 13 (0.011 s^−1^μM^−1^ vs 0.0036 s^−1^μM^−1^), suggested that its intracellular abundance could be low due to degradation by cellular proteases (38, 61). We confirmed this using Western blots with anti-DHFR polyclonal antibodies (Fig. 7B and Fig. S10 in the Supporting Material). No DHFR-specific band was detected in the soluble fraction of lysates from the strain carrying the Chimera 1 gene, whereas such bands were visible for the remaining chimeric strains, albeit less intense than that of the wild type. As in previous studies (33), the intracellular abundance determined for the tested chimeras follows a general inverse correlation with the fluorescence intensity upon *bis*-ANS binding to the purified proteins (Fig. S11 in the Supporting Material). The *bis*-ANS dye binds preferentially to hydrophobic regions exposed in molten globule conformations. These results are compatible with the non-native α_B_-helix conformations observed in simulations of Chimeras 1 and 6, which may increase the likelihood of molten globule states. The Chimera 19 strain, however, also has a lower intracellular abundance of DHFR despite native-like simulation conformations, which points to aspects of DHFR conformation and cellular interactions either not captured or not accurately rendered by simulations.

We reasoned that enhancing the intracellular protein abundance of strains with Chimeras 1 and 6 could rescue their growth. We engineered versions of these chimeric strains with a promoter mutation (C-35T) that is known to increase the expression of wildtype DHFR several-fold (62). As expected, the intracellular abundances of Chimeras 1 and 6 increased considerably on the background of the *folA* promoter mutation (Fig. 7C), and both strains were able to grow without metabolic supplementation (Fig. 7D). The fitness of the chimeric strains is in good agreement with the predictions made by a DHFR fitness model we described previously (38) that is based on intracellular abundance and biophysical properties of the variants (Fig. 7E). This model predicts that none of the other non-orthologous chimeras studied in this work would result in a strain that can grow without folAmix supplementation. These results show that the replacement of the α_B_-helix in DHFR with non-orthologous sequences has a strongly deleterious effect on fitness that is due to decreases in catalytic function and stability against intracellular degradation. Nevertheless, the viability of strains with Chimeras 13 and 19 shows that certain, sequence-compatible α-helix swaps are possible, leading to solutions that are functional in the biological context.

## Discussion

The decomposition of protein structure into discrete segments has been a productive method in protein structural analysis due to structural degeneracy in sequence space (63, 64). Sub-domain modularity may be a fundamental property of proteins that facilitates their evolution by promoting robustness and evolvability (7, 16, 65). We define modules as independent and transferable protein segments that maintain consistent structure despite changes in structural context. We examine the extent to which this kind of modularity is present in α-helices using the DHFR α_B_-helix. The α_B_-helix of DHFR has several typical characteristics seen in helix-sheet structures, including alignment of the helix axis with β-strands and the presence of hydrophobic aromatic residues on the helix and hydrophobic aliphatic residues on the β-sheet (66). Studying this helix provides a highly sensitive way to probe the effects of helix replacements since conformational changes that affect the positioning of the D27 sidechain are expected to impact catalytic activity considerably.

Substitution of an α-helix for structurally similar sequences does not preserve the wildtype-like helix conformation in most cases, with pleiotropic and deleterious effects on thermal stability, catalytic activity, and cellular abundance, which together determine cellular fitness (38, 61). We find principles that govern the outcome of whole-segment substitutions, namely biophysical compatibility, as characterized by EVcouplings and MD simulations. Evolutionary coupling analysis has been shown to predict the experimentally-measured fitness effects of mutations (44). Here, we show that EVcouplings can also predict the effects of larger scale sequence changes on enzyme catalysis, which partly determines fitness. In our study, the EVcouplings non-epistatic model Δ*E* correlates slightly better with *in vitro* properties than Δ*E* calculated by the epistatic model. This may be because simultaneously mutating several residues in the α_B_-helix puts chimeric DHFRs in less populated sequence space where epistatic coupling parameters are less certain.

Our MD simulations appear to be in line with past results in which computational techniques predict thermal stability more easily than catalytic activity (67, 68). The finding that increased hydrophobic surface area of helix residues strongly correlates with increased thermal stability is curious. The general expectation is that introduction of hydrophobic surface residues is destabilizing (69), but there are examples in which the introduction of hydrophobic surface residues stabilizes a protein (70–72). Here, changes in helix conformation accompany changes in hydrophobic surface area, and it is difficult to identify a mechanism relating changes hydrophobic surface area to thermal stability. A past study in our group characterized destabilizing mutations in DHFR and found that such mutants formed soluble oligomers at 42 °C that prevented aggregation (73). Variation in hydrophobic surface area may relate to oligomerization of DHFR at increasing temperatures, which would be expected to protect against thermal denaturation.

Of the non-orthologous chimeras, Chimera 19 is particularly notable in that its sequence identity with *E. coli* DHFR is low and that the Chimera 19 *E. coli* strain is viable, demonstrating that in some cases, the DHFR α_B_-helix can be switched for an evolutionarily unrelated sequence. On the other hand, the low catalytic activity and non-wildtype conformations in most non-orthologous chimeras demonstrate incompatibility. One key issue with low activity chimeras appears to be conformations that occlude the binding site. This suggests that one constraint on the α_B_-helix sequence is avoidance of errant interactions with the hydrophobic DHF binding pocket next to the helix. Chimera 1 is a major exception to the trends. The numerous charged sidechains in the Chimera 1 α_B_-helix lead to flexible non-helical conformations that do not block the DHF binding site. The binding of DHF may be sufficient to induce transient folding of the α_B_-helix to catalyze the enzymatic reaction, or it may even be possible that the other negatively charged residues in the Chimera 1 replacement sequence promotes DHF reduction (74). Structural instability comes at the cost of intracellular abundance, however, and increased expression levels are necessary for growth of the Chimera 1 *E. coli* strain. Thus, although the disordered Chimera 1 α_B_-helix is not a sequence that that retains helicity in a new context, Chimera 1 demonstrates an alternative solution for catalytic activity.

The results of α_B_-helix substitution in this work can be compared with previously characterized point mutations in the DHFR α_B_-helix. Such point mutations have much smaller effects on stability and catalysis compared to helix substitution. For instance, L28R and A26T are trimethoprim resistance mutations that result in a 50% decrease in *k_cat_*/*K_M_*, with L28R improving thermal stability (38, 75, 76). Single mutants L28F and L28Y are also within an order of magnitude of wildtype *k_cat_*/*K_M_* (77). In this work, we identify W30 and F31 as key residues for maintaining enzyme function and wildtype-like helix conformations, since the chimeras with the highest activity retain at least one of W30 and F31. Single mutants W30G and W30R are comparable in catalytic activity to wildtype DHFR (75). Mutations at position 31 are more deleterious however. F31Y, F31V, and L28A/F31A have been characterized, with the latter two resulting in two orders of magnitude loss in *k_cat_*/*K_M_* (78–80). All but one of the non-orthologous chimeras are mutated at position 31, and the combined effect of multiple mutations over the entire helix produces a severe loss of catalytic activity.

One framework with which to interpret our results is fold polarity (81, 82). This hypothesis states that a protein fold with higher polarity—a fold characterized by both a well-ordered scaffold and an active site with flexible loops—is simultaneously robust to mutations and also more capable of evolving new functions. *E. coli* DHFR was cited as an example of a low polarity fold since the DHF binding site in DHFR is composed primarily of residues in β-sheets and α-helices (81). This framework suggests that the DHF binding site in DHFR cannot easily adjust to mutations compared to proteins with more polar folds. It is possible that whole-helix substitutions in locations further from the active site or in proteins with greater fold polarity would be less deleterious to protein function, as was seen with *E. coli* alkaline phosphatase (30). Our results show that catalysis is more sensitive to helix substitution than thermal stability: although the *k_cat_*/*K_M_* values of non-orthologous chimeras are uniformly worse than those of orthologous chimeras, the *T_m_* values of several non-orthologous chimeras are comparable to some of the orthologous chimeras. Furthermore, preserving helical secondary structure upon switching an α-helix is more easily achieved than preserving catalytic activity. MD simulations of non-orthologous chimeras find seven variants that maintain high helicity, and EVcouplings analysis shows that the helical sequences in this study are slightly more compatible replacements for the α_B_-helix than are random subsequences taken from the entire *E. coli* proteome.

## Conclusion

By testing the fungibility of α-helices, we investigated the principles governing α-helical conformations in proteins: maintenance of interactions by key residues and avoidance of errant interactions. Our results highlight how local destabilization caused by the non-compatibility of the replacement sequences, which leads to low catalytic function and higher intracellular protein degradation, places limits on helix modularity. Our work reveals the relationship between broad sequence variation and molecular and fitness effects for an essential enzyme that is central to key metabolic pathways.

## Author contributions

**Victor Y. Zhao**: Investigation, Methodology, Software, Visualization, Writing – original draft. **Joao V. Rodrigues**: Investigation, Methodology, Visualization, Writing – original draft. **Elena R. Lozovsky**: Investigation, Methodology. **Daniel L. Hartl**: Conceptualization, Supervision, Writing - Review & Editing. **Eugene I. Shakhnovich**: Conceptualization, Funding acquisition, Supervision, Writing - Review & Editing.

## Acknowledgments

This work was supported by the National Institute of General Medical Sciences of the National Institutes of Health (RO1GM068670) and the National Science Foundation Graduate Research Fellowship Program (DGE1745303, awarded to V.Y.Z.). The computations in this paper were run on the FASRC Cannon cluster supported by the FAS Division of Science Research Computing Group at Harvard University. This work also used the Extreme Science and Engineering Discovery Environment (XSEDE), which is supported by National Science Foundation grant number ACI-1548562 (83). Specifically, this work used the Bridges cluster at the Pittsburg Supercomputing Center and the Comet cluster at the San Diego Supercomputer Center through allocation BIO200028. Plots were created using the Seaborn and Matplotlib Python packages (84, 85). We are grateful to Rostam M. Razban for a critical reading of the paper.

## Supporting References

References (86–90) appear in the Supporting Material.

## Notes

### Competing Interest Statement

The authors have declared no competing interest.

### Summary of Updates

Revised / streamlined narrative.

