## supplementary_information for "Switching an active site helix in dihydrofolate reductase reveals limits to sub-domain modularity"

### Extended Methods

#### DHFR chimeric constructs

To be able to carry out *in vitro* characterization of a considerable number of chimeras, we first developed an efficient two-step PCR strategy to generate various DNA constructs and clone them into an expression plasmid. DNA constructs of chimeric DHFR fused to a C-terminal (6x) His-tag were generated and cloned into the plasmid pFLAG-CTC by a series of PCR reactions using as template the plasmid pFLAG-CTC-wtDHFR, in which wild type E. coli DHFR had been cloned using NdeI/HindIII restriction sites (1). First, separate fragments of ~550 bp size that flank the helix segment to be modified, designated pFLAG_A and pFLAG_B, were amplified using the set of primers Ampl_pFLAG_H1/ H1_pKD_rev and H1_pKD_for/Ampl_pFLAG_H1_rev, respectively. The backbone of pFLAG-CTC-wtDHFR was amplified using H1_downstream_for/ H1_upstream_rev to produce a linear product (TpFB) where the sequence of the wild type helix segment is absent. Second, for each helix sequence, we designed pairs of complementary primers containing the helix-specific sequence and fixed overhangs with homology either end flanking the helix segment. These pairs of primers were combined with fragments pFLAG_A and pFLAG_B, and a ~1kb linear fragment was amplified by PCR using Ampl_pFLAG_H1/ Ampl_pFLAG_H1_rev. Finally, the amplified product, containing the desired chimeric DHFR sequence, was used to extend the fragment TpB by circular polymerase extension using Phusion polymerase. The non-ligated circular product was used to transform DH5alpha cells, and the sequence of the transformants was verified by Sanger sequencing. For a single colony being sequenced per each chimeric construct, an approximately 75% success rate was achieved with this methodology. See Table S2 for a list of primers used in this work.

#### *bis*-ANS fluorescence

Purified chimeric proteins (2 µM) were incubated for 5 minutes at 37˚C in the presence of 12 µM *bis*-ANS, in 20 mM potassium phosphate buffer pH7.2. The fluorescence emission spectra were then recorded between 400 and 600 nm (slit 5 nm) upon excitation at 395 nm (slit 2.5 nm). The emission band was integrated, and the fluorescence of *bis*-ANS alone was subtracted. Results are normalized with respect to the wild type protein and represent the average of 3 measurements.

#### Size exclusion chromatography

A Superdex 10/300 GL analytical column (Cytiva) was equilibrated with 20mM tris, 150 mM NaCl, pH 8.0 at a flow rate of 0.8 mL/min. 75 µL of a solution of each protein (20 µM) and NADPH (100 µM) were sequentially injected using an Akta 905 Autosampler. As molecular standards, we used a mixture containing bovine serum albumin (66 kDa), carbonic anhydrase (29 kDa), cytochrome c (12.3 kDa), and aprotinin (6.5 kDa).

#### Agadir score

To predict the helical content of the sequences that were used to replace the α-helix in DHFR, we used the Agadir score (2–6), which was computed based on the 10-residue (positions 25-34) subsequence corresponding to each replacement helix. The parameters used in the calculations were: temperature = 37˚C, ionic strength = 0.1 M, pH = 7.0.

#### MD simulation structure preparation and settings

Structure preparation began with an X-ray crystal structure of E. coli DHFR with NADP+ and folic acid bound (PDB ID 4NX6). MOE (7) was used to perform structure preparation, with the PFROSST forcefield loaded. The crystallographic waters, manganese ion, and beta-mercaptoethanol molecule were deleted. The Structure

Preparation tool was used to optimize side chain protonation states. His 45 and 114 were protonated at the ε nitrogen, His 124 and 149 were protonated at the δ nitrogen, and His 141 was doubly protonated. From this prepared structure, all DHFR chimeras were prepared using the Homology Model tool. FASTA sequences of chimeric DHFRs were loaded, and the prepared DHFR complex was used as the template for homology modeling, with ligand atoms set as part of the molecular environment. The NADP+ and folate ligands were then removed to obtain the apo structures of each DHFR variant.

Proteins structures were solvated with water in cubic boxes with at least a 10 Å distance between box edges and protein atoms. Na+ and Cl- ions were added to the system at a concentration of 150 mM such that the system charge was neutralized. The AMBER ff14SB forcefield was used to model protein atoms, and the TIP3P forcefield was used for the water (8, 9). All MD simulations were performed using NAMD version 2.14 (10), and the included REST2 script was used for replica exchange simulations. All atoms in protein residues 24 through 36 were tempered. The nonbonded interaction cutoff distance was set to 9 Å; the simulation timestep was 2 fs, and all bonds to hydrogens were constrained. Long-range electrostatics were calculated using particle-mesh Ewald with a grid spacing of 1.0 Å. Simulations were run in the NPT ensemble at 300 K and 1 atm. The temperature was maintained by a Langevin thermostat with a damping value of 1 $ps^{-1}$, and the pressure was maintained by a Langevin piston with a period of 100 fs and decay of 50 fs. Simulation coordinates were saved every 0.5 ps.

Initial equilibration was performed by minimizing the system for 5000 steps and then running 5 ns of simulation. Simulation coordinates from the end of the equilibration trajectory were used to initiate replica exchange simulations. Replica exchange simulations were run for 300 ns, with some simulations extended to 375 ns to achieve converged helix RMSD values in the last 100 ns of simulations (Supplementary Fig. S5). The last 100 ns of simulations of the non-scaled replicas ($\beta_m = \beta_0$) were used for measurements.

#### Simulation analysis

Simulation trajectories were analyzed using MDTraj version 1.9.4 (11). The *E. coli* wildtype DHFR crystal structure (PDB ID 4NX6) was used as the reference for RMSD measurements, and the following are the residue selections: adenosine-binding subdomain: 38-88; adenosine-binding subdomain core: 39-43, 58-62, 73-85; major subdomain: 1-37, 89-159; major subdomain core: 2-8, 91-93, 109-115, 133-141, 151-158; $\alpha_B$-helix: 25-35; Met 20 loop: 9-24; FG loop: 116-132; GH loop: 142-150.

For all RMSD measurements except for RMSD of the entire adenosine-binding and major subdomains, measurements were performed by first aligning simulation structures to the crystal structure reference using the Cα-atoms of core residues of the major subdomain. Then, RMSD measurements were performed over the Cα-atoms of the selected residues of interest. Helix helicity was calculated by DSSP (12).

The Shrake-Rupley method (13) within MDTraj was used to measure the solvent accessible surface area (SASA) with default parameters (probe radius 1.4 Å and 960 sphere points). This measures the SASA of each atom in a protein structure. A simple heuristic, following Shrake and Rupley, was used to classify protein atoms as either non-polar or polar: A hydrogen atom is nonpolar only if it is bonded to carbon. A carbon atom is nonpolar except when it is bonded to two or more heteroatoms. Sulfur atoms are nonpolar. All other atoms are polar. The SASA values of non-polar atoms of helix atoms (residues 25-35) were summed to determine the hydrophobic SASA of the helix.

Binding site occlusion was calculated as follows: For each simulation frame, the protein coordinates were aligned to the *E. coli* DHFR crystal structure (PDB ID 4NX6). The coordinates of the folate ligand from the crystal structure were then added to the simulation frame. A 3D grid with a spacing of 0.1 Å was overlaid in the vicinity of the ligand. The volume available to the folate ligand was then measured by the number of grid points that are within the van der Waals radius of any ligand atom but not within the van der Waals radius of any protein atom. Occlusion was calculated as the fraction of ligand volume blocked by protein atoms.

### Supplementary figures and tables

Table S1: List DHFR chimera sequences and properties. Available as separate Excel spreadsheet.

Table S2: Primers used in this work.

| | | |
|---|---|---|
| pFLAG Upstream Fragment (pFLAG_A) | Ampl_pFLAG_H1_for CCGGGAGCTGCATGTGTCAGAGG | H1_pKD_rev CAGGTTCCACGGCATGG |
| pFLAG Downstream Fragment (pFLAG_B) | H1_pKD_for ACCTTAAATAAACCCGTGATTATGG | Ampl_pFLAG_H1_rev AATTCTGTTTTATCAGACCGCTTCT |
| Truncated pFLAG Backbone (TpFB) | H1_downstream_for GCCGCCATACCTGGGAATCAATC | H1_upstream_rev ACGCCGCAATCAGACTGATCATATG |
| pKD13.2 Upstream Fragment (pKD13.2_A) | Ampl_RRfolA_for GTGCCGATCAACGTCTCATTTTCG | H1_pKD_rev |
| pKD13.2 Downstream Fragment (pKD13.2_B) | H1_pKD_for | Ampl_RRfolA_rev GCTTCCTCGTGCTTTACGGTATCG |
| Truncated pKD13.2 Backbone (TpKD) | H1_downstream_for | H1_upstream_rev |
| Helix-primers | 30nt-forward-helix-sequence $(NNN)_2$-GAT-$(NNN)_7$-ACCTTAAATAAACCCGTGATTATGG | 30nt-reverse-helix-sequence $(NNN)_7$-CTA-$(NNN)_2$-CAGGTTCCACGGCATGGCGTTTT |
| Amplification of linear PCR for gene editing | PCRseq_KefC_for2 CTGCTCGGTTTCCTCATCATCAA | PCRseq_apaH_rev CGTCCCTTTCAGCATCGACATT |
| Sequencing | PCRseq_RRfolA_rev GCCTTCTATCGCCTTCTTGACGA | |

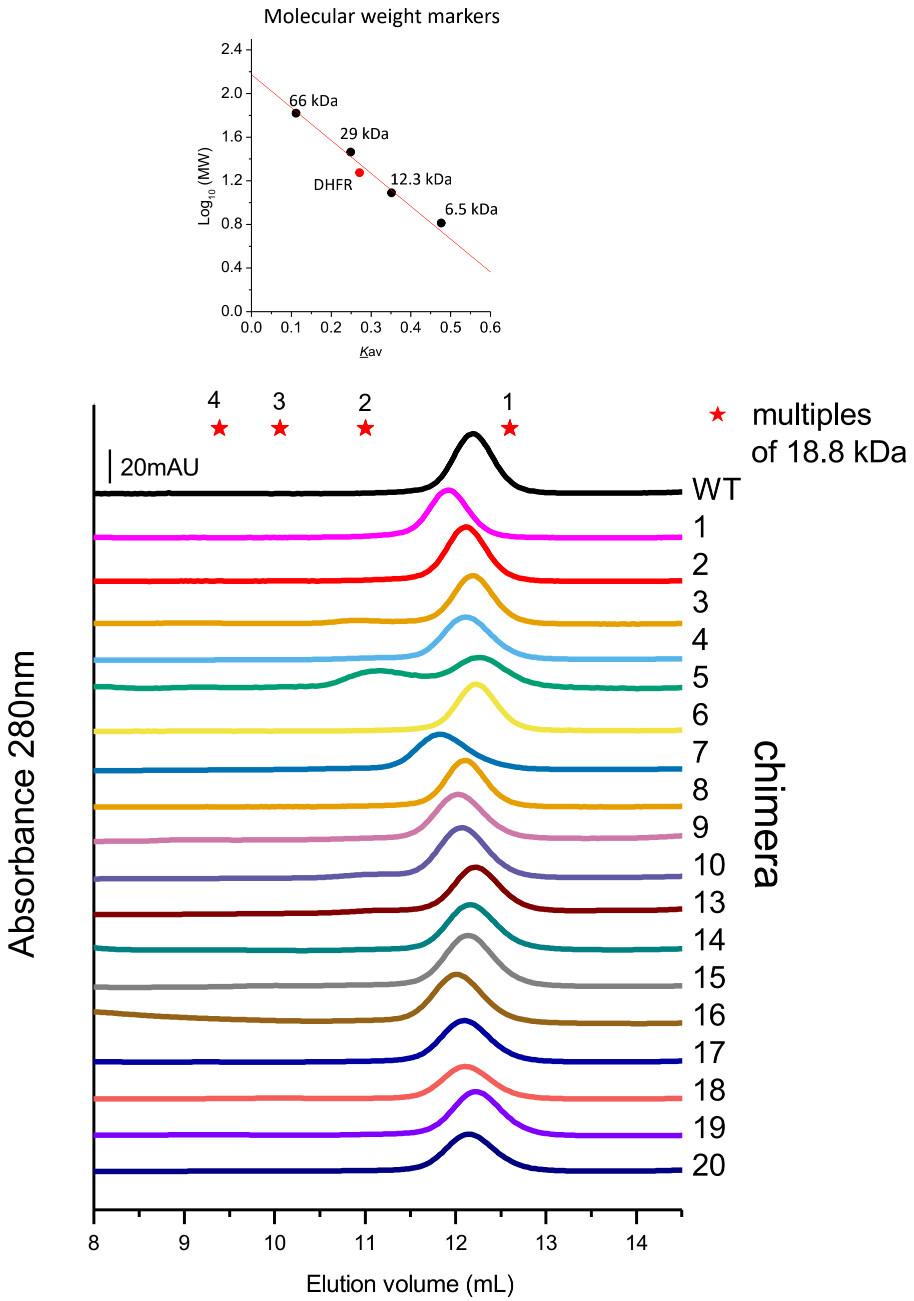


Fig. S1: Size exclusion analysis of purified chimeras. Chimeric DHFR are monomeric, except for chimera 5. Size exclusion chromatograms on a Superdex 75 GL300/10 of purified DHFR shows that the wild type protein elutes with an apparent molecular weight of 22.6 kDa. Chimera 5 elutes both as a dimer and monomer.

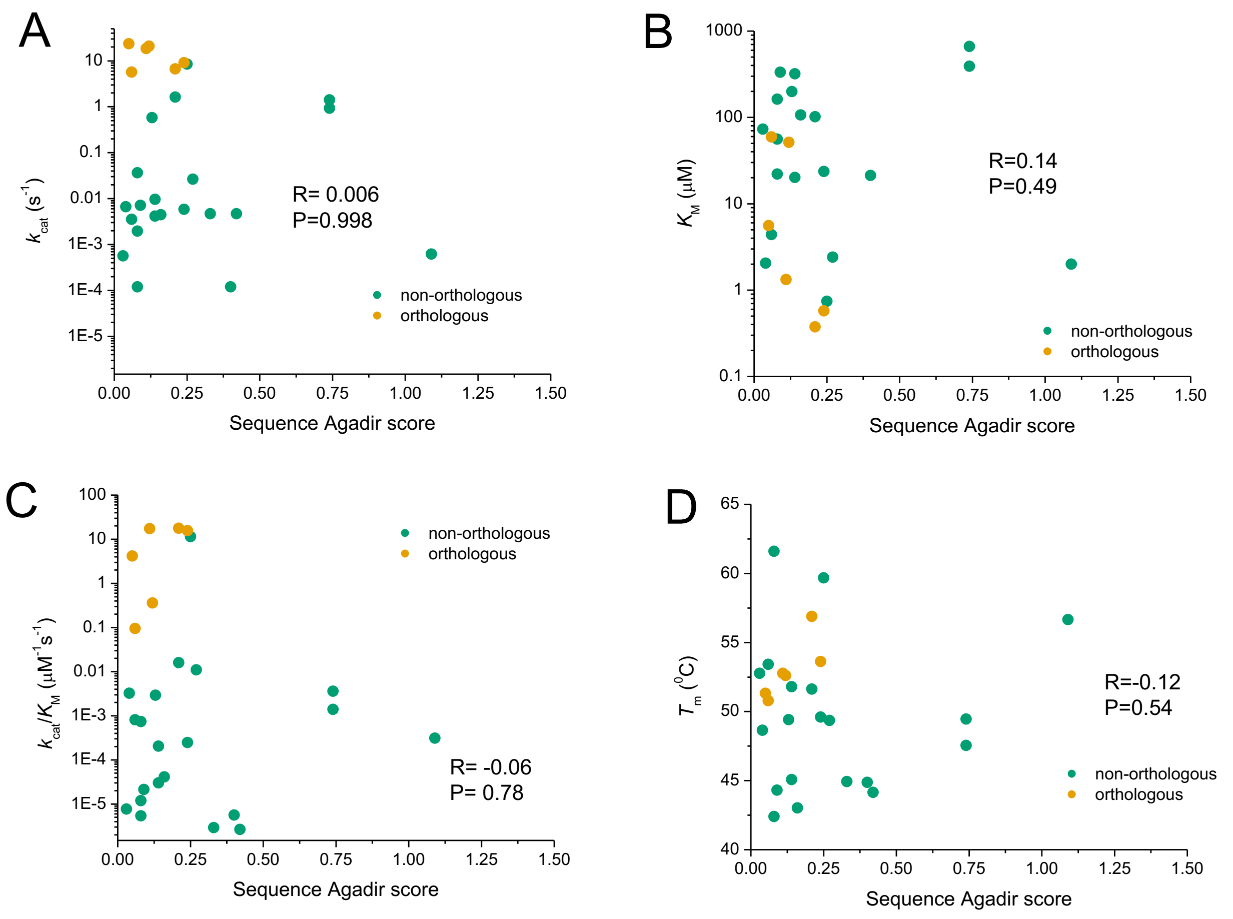


Fig. S2: *In vitro* properties of chimeric DHFRs do not correlate with the intrinsic helical propensity of the sequences. Agadir score predicts the helical propensity of isolated peptide sequences (see Experimental procedures). Spearman rank correlation coefficients between *in vitro* properties and sequence Agadir scores were calculated using data for all DHFR chimeras (orthologous and non-orthologous).

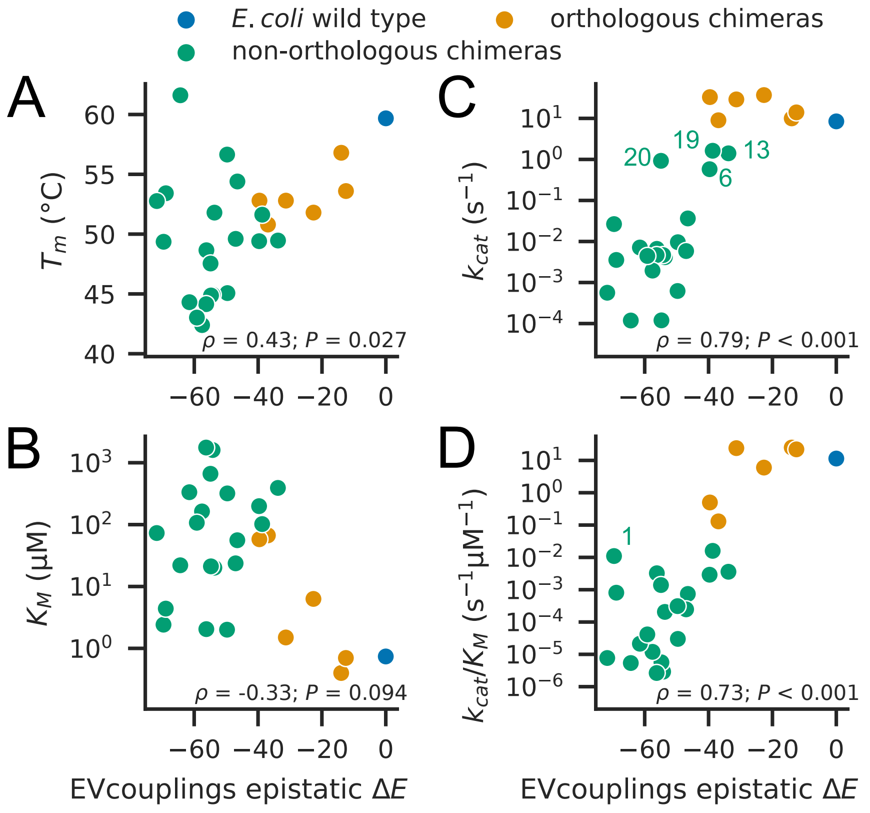


Fig. S3: EVcouplings epistatic model Δ*E* correlates with several measured *in vitro* properties of DHFR chimeras, with slightly lower correlation coefficients compared to the non-epistatic model. Epistatic model Δ*E* is compared with (A) $T_m$, (B) $K_M$, (C) $k_{cat}$, with Chimeras 6, 13, 19, and 20 labeled, and (D) $k_{cat}/K_M$, with Chimera 1 labeled. Spearman rank correlation coefficients and *P* values are given in the lower portion of each plot.

Table S3: Spearman correlation coefficients (*P* values in parentheses) between chimera sequence measures and *in vitro* properties. Sequence similarity was calculated according to Grantham's distance between amino acids (14).

| | $T_m$ | $k_{cat}$ | $K_M$ | $k_{cat}/K_M$ |
|---|---|---|---|---|
| EVcouplings epistatic model $\Delta E$ | 0.41 (0.04) | 0.78 (< 0.001) | -0.30 (0.13) | 0.73 (< 0.0001) |
| EVcouplings independent model $\Delta E$ | 0.37 (0.064) | 0.84 (< 0.001) | -0.27 (0.18) | 0.74 (< 0.0001) |
| Sequence identity | 0.27 (0.19) | 0.63 (< 0.001) | -0.24 (0.24) | 0.57 (0.002) |
| Sequence similarity (Grantham) | 0.27 (0.18) | 0.63 (< 0.001) | -0.24 (0.22) | 0.58 (0.0014) |

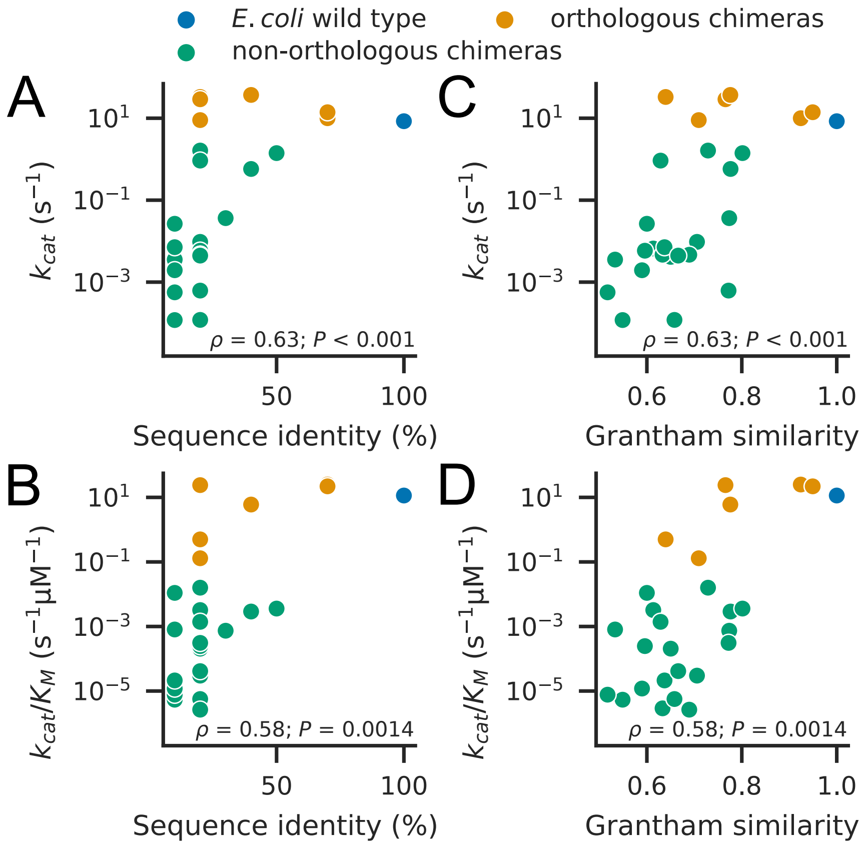


Fig. S4: Sequence identity and sequence similarity correlate with $k_{cat}$ and $k_{cat}/K_M$, although the prevalence of variants with 10% or 20% identity makes sequence identity less informative. Sequence identity is compared with (A) $k_{cat}$ and (B) $k_{cat}/K_M$. Grantham similarity is compared with (C) $k_{cat}$ and (D) $k_{cat}/K_M$. Grantham similarity scores are scaled such that wildtype DHFR has a score of 1. Spearman rank correlation coefficients and $P$ values are given in the lower portion of each plot.

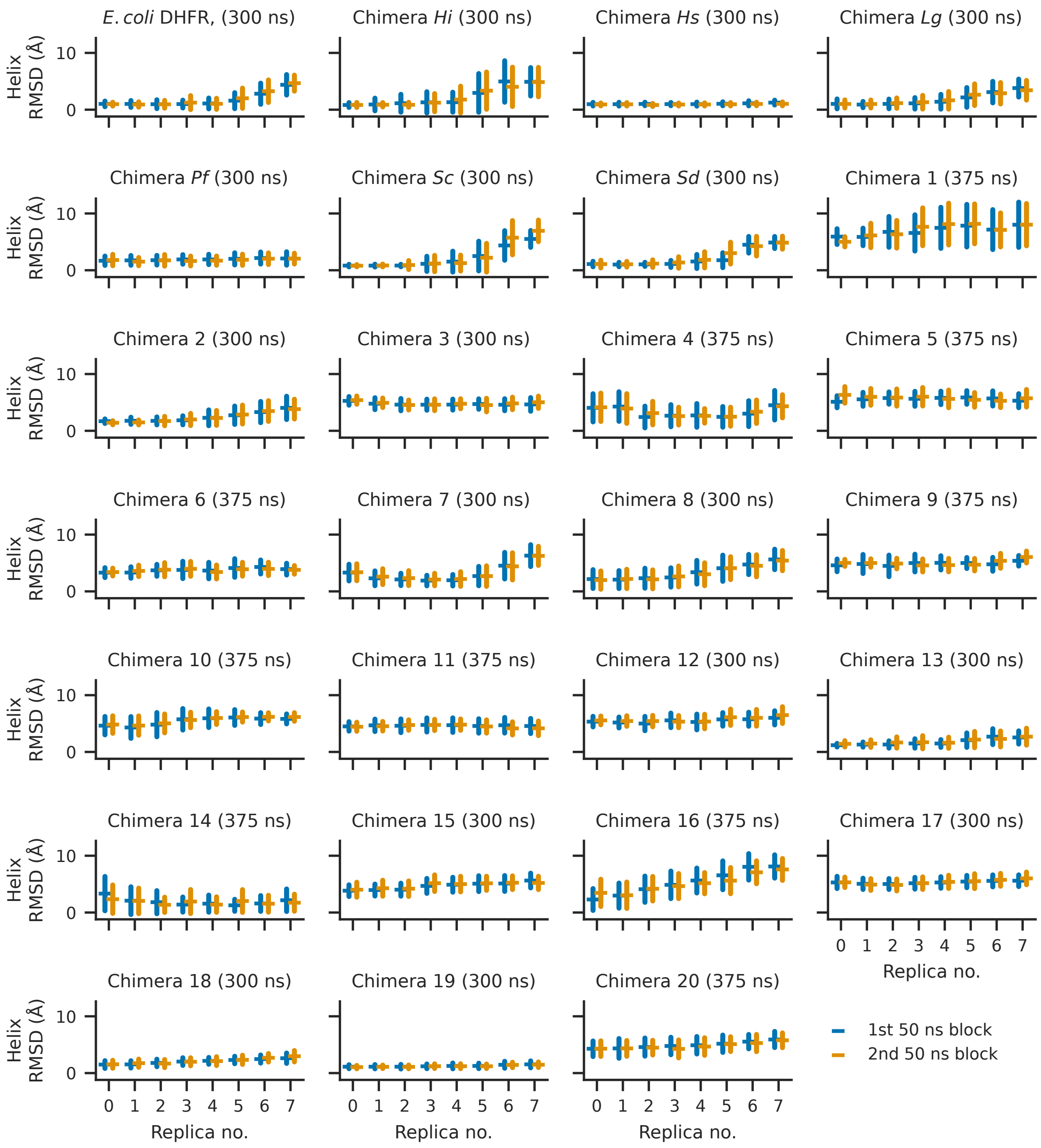


Fig. S5: Convergence of simulation trajectories. Throughout this work, the last 100 ns of replica exchange simulations for each chimera were used for analysis. Here, $\alpha_B$-helix RMSD over the first and second halves of the 100 ns window are compared. Replica 0 corresponds to the physically-relevant, full-potential state. Simulation length for each DHFR variant is indicated in parentheses. Error bars indicate standard deviations.

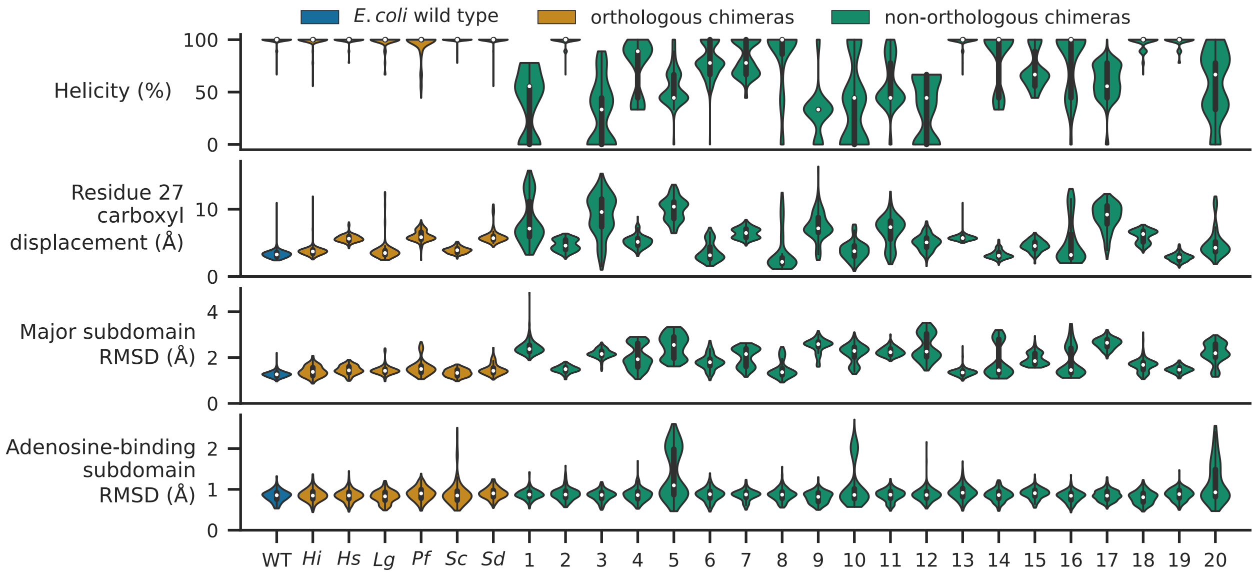


Fig. S6: Distributions of simulation observables by chimera. Residue 27 carboxyl displacement measures the RMSD of the residue 27 sidechain carboxyl centroid, after alignment of major subdomain core residues. Note that Chimeras *Hs*, *Sc*, and *Sd* have E substituted for D at position 27. Major subdomain RMSD and adenosine-binding subdomain RMSD represent RMSD measurements of the two subdomains of DHFR. Inner white marker denotes median value, and ends of thicker vertical bars denote 25$^{th}$ and 75$^{th}$ percentiles. (For Chimera 9, 25$^{th}$ and 75$^{th}$ percentile values for helicity match the median value of 33%.)

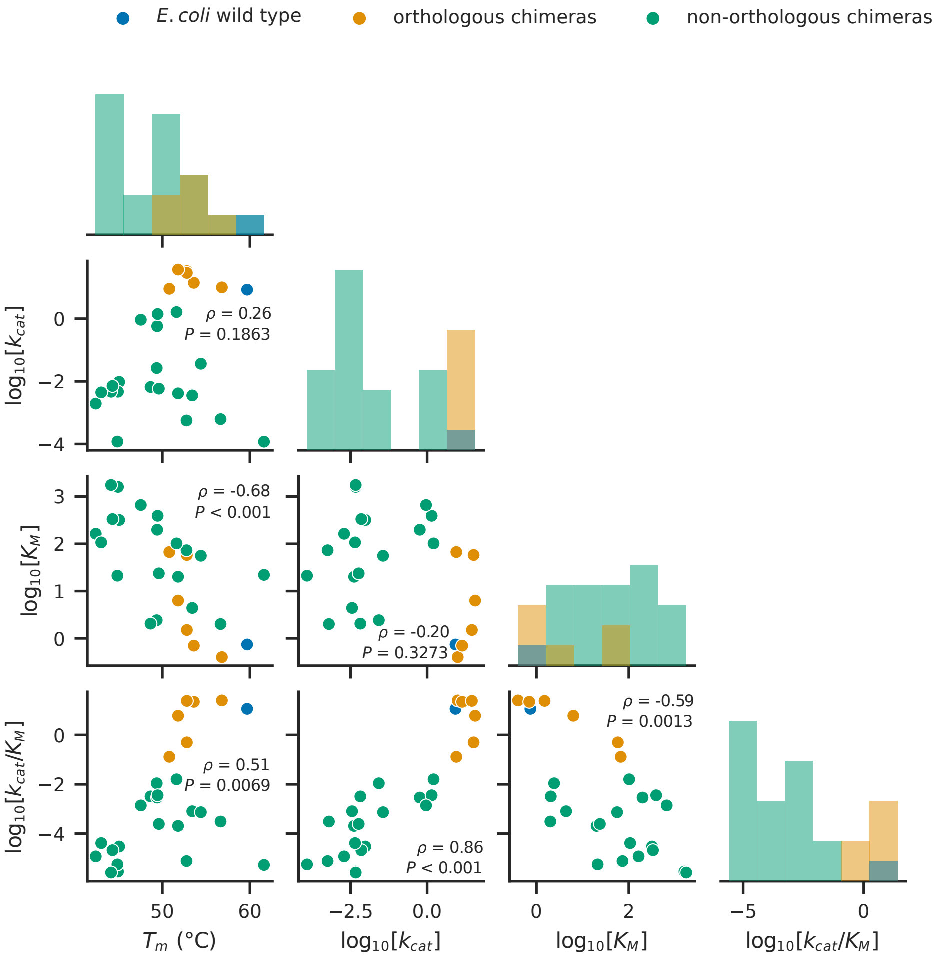


Fig. S7: Correlation between key experimental *in vitro* properties. Spearman rank correlation coefficients and *P* values are given in each plot.

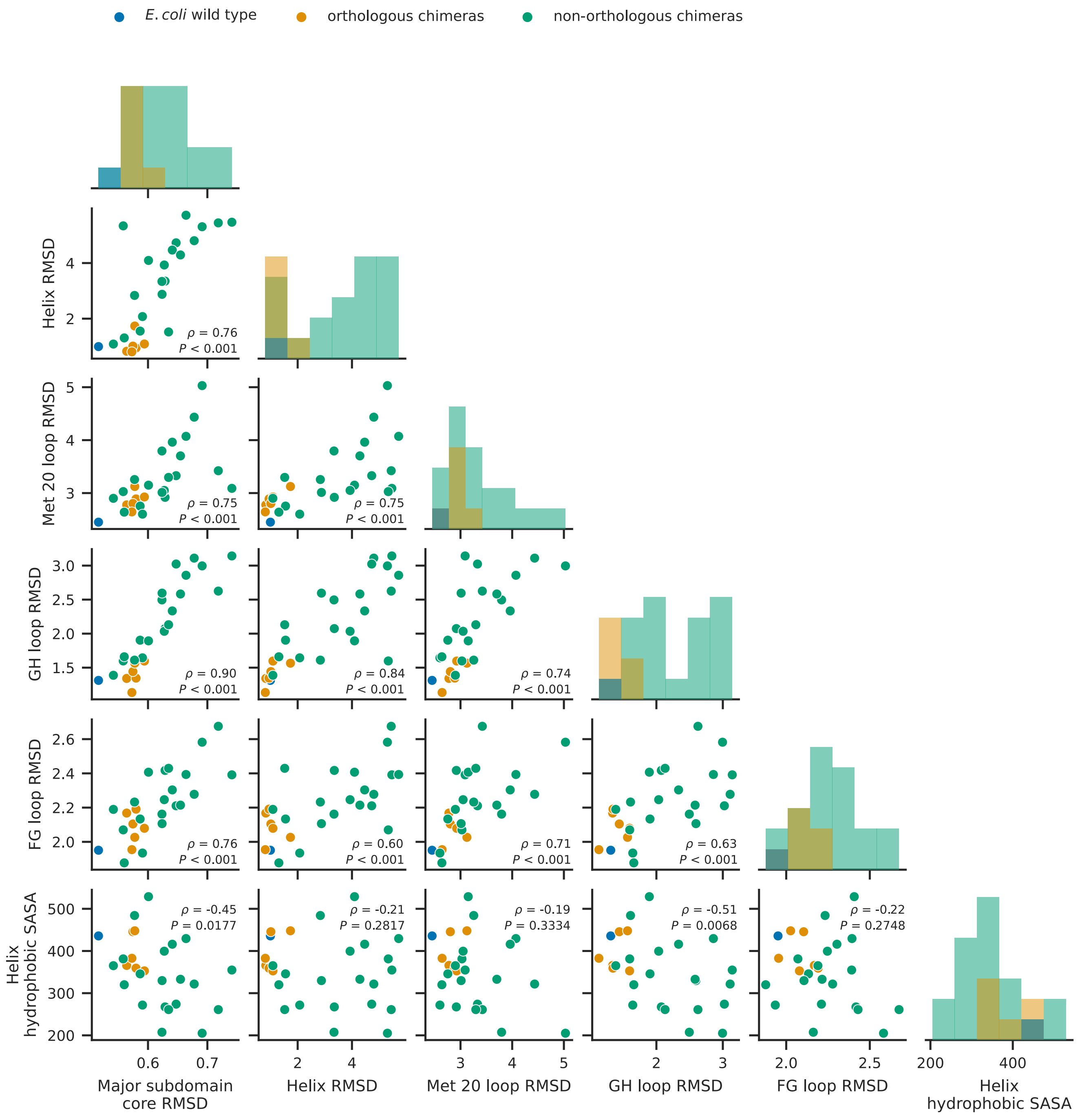


Fig. S8: Most simulation order parameters are correlated with each other. Helix hydrophobic SASA is less correlated with RMSD measurements. Scatterplots showing correlations between key simulation order parameters are plotted. Spearman rank correlation coefficients and *P* values are given in each plot. All RMSD measurements use angstroms; units of SASA are square angstroms.

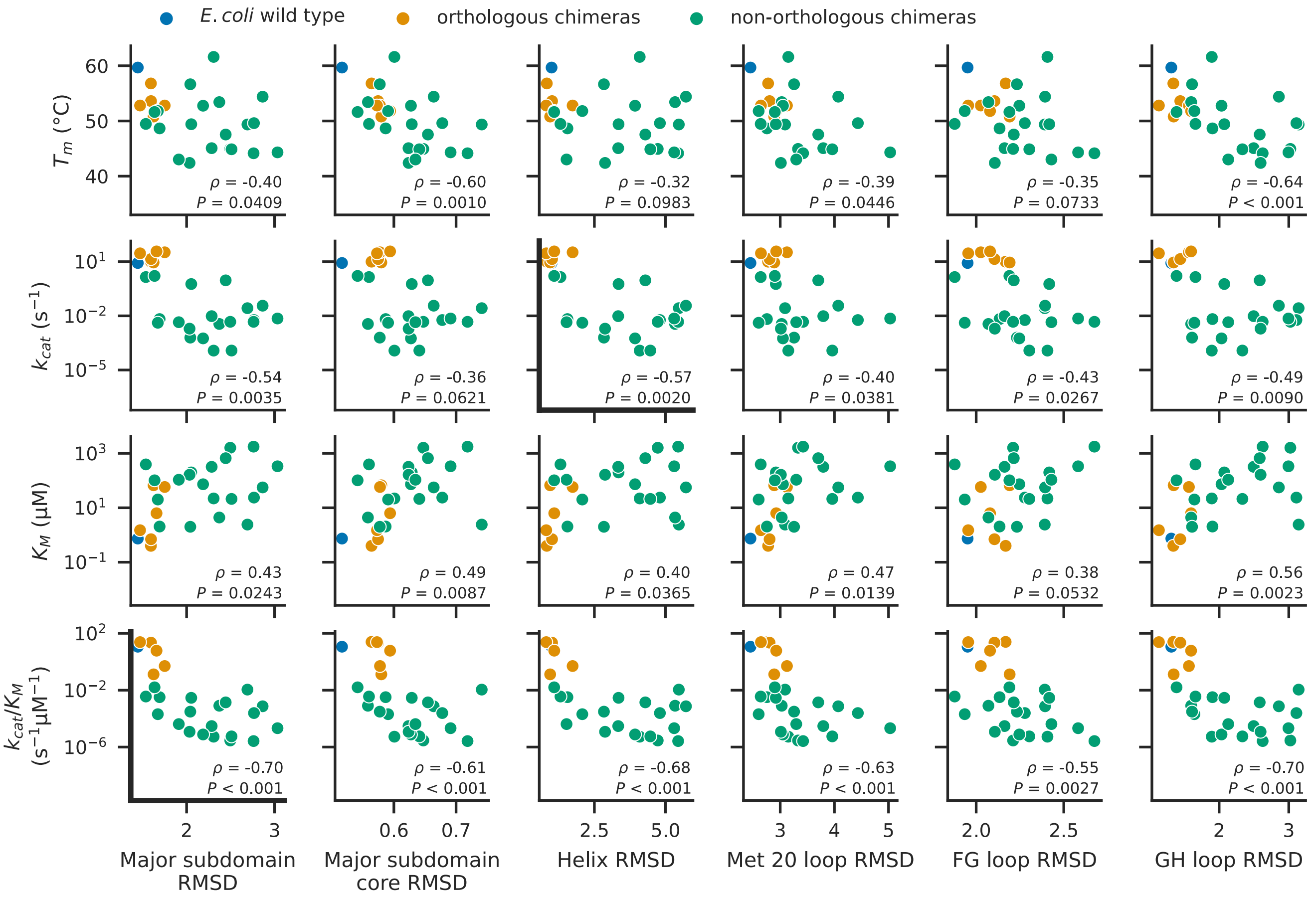


Fig. S9: Simulation order parameters differentiate between orthologous and non-orthologous chimeras. Scatterplots of simulation mean RMSD for different portions of the major subdomain versus *in vitro* measurements of DHFR variants. Bold axes denote the two subfigures shown in Fig. 5. Spearman rank correlation coefficients and *P* values are indicated in the bottom right of each plot. When only non-orthologous chimeras are considered, only the relationship between GH loop RMSD and $T_m$ retains statistical significance ($\rho$ = -0.51, $P$ = 0.023). Chimera 1 is indicated in several plots. All RMSD measurements use angstroms.

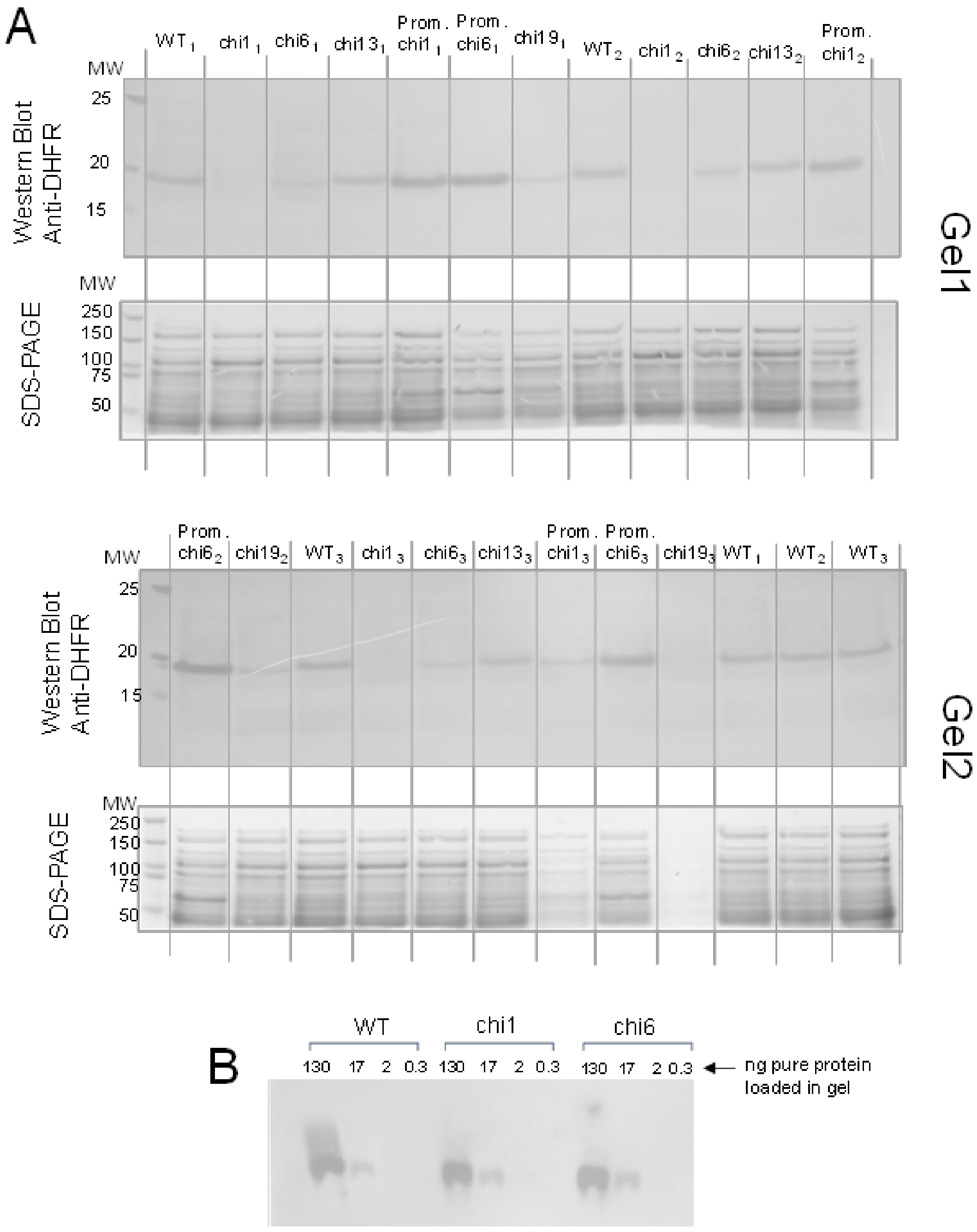


Fig. S10: Western blot analysis to quantify the intracellular abundance of chimeric DHFR proteins in chromosomal replacement strains. (A) Soluble fraction of lysates from different chimeras was loaded on an SDS-PAGE gel. *The* low-MW *half of the gel* was transferred to a nitrocellulose membrane for Western blotting to detect DHFR-specific band using polyclonal anti-DHFR antibodies. The high-MW half of the gel was directly stained with Coomassie and used to *control for* the total protein content that was loaded in the gel. (B) Polyclonal anti-DHFR antibodies react similarly against both the wild type and chimeric proteins.

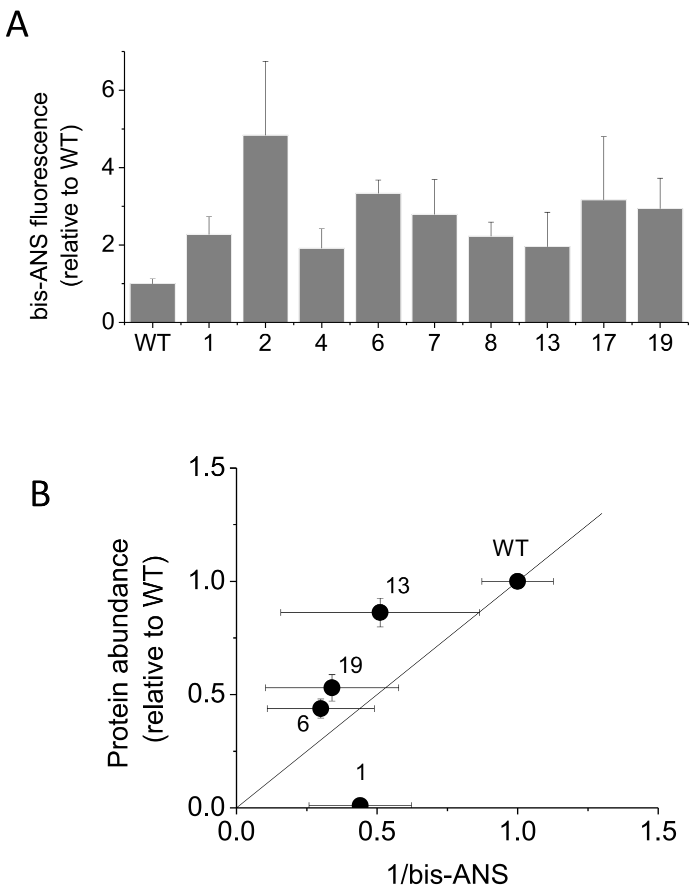


Fig. S11: Increased bis-ANS binding to chimeric DHFR proteins indicates higher degree of molten-globule conformations with respect to the wild type protein. (A) Bis-ANS (12 µM) was mixed with purified chimeric DHFR poteins (2µM) in 20 mM potassium phosphate buffer, pH 7.2 and the solution was incubated for 5 min at 37˚C. The fluorescence emission spectra (excitation 395nm) were then measured between 400 and 600 nm. The intensity of the emission band was integrated, the background bis-ANS fluorescence subtracted, and the values were normalized to that of the wild type protein. (B) The reciprocal of bis-ANS fluorescence intensity measured for purified chimeras predicts the intracellular abundance of chimeric DHFR in strains with chromosomal replacement of the *folA* gene.
